# FGR Src Family Kinase Causes Signaling and Phenotypic Shift Mimicking Retinoic Acid-Induced Differentiation of Leukemic Cells

**DOI:** 10.1101/2024.08.21.608654

**Authors:** Noor Kazim, Wang Peng, Jianbo Yue, Andrew Yen

## Abstract

Retinoic acid (RA) is an embryonic morphogen used in cancer differentiation-therapy. It causes a plethora of changes in gene expression culminating in cell differentiation. We now find that amongst them, expression of the Src-family-kinase, FGR, by itself causes cell differentiation analogous to RA. The historically dominant/classical paradigm for RA mechanism of action is transcriptional activation via binding to the ligand-activated nuclear receptors, RAR/RXR. In the HL-60 human myelo-monocytic leukemia model, an actively proliferating, phenotypically immature, lineage bi-potent NCI-60 cell line, RA causes election of the myeloid lineage and phenotypic maturation with G1/0 growth inhibition. It thereby converts transformed immature proliferating tumor cells to mature growth retarded cells that bear fidelity to non-transformed mature myeloid cells. The present study finds that expression of the FGR SFK(SRC-family-kinase) alone is sufficient to induce differentiation. Akin to RA, the phenotypic conversion manifests as expression of CD38, CD11b, and ROS, as well as the p27(kip1) CDKI (cyclin-dependent-kinase-inhibitor that retards cells in G1/0) characteristic of mature myeloid cells. To pursue mechanistic insight, signaling attributes known to promote RA-induced differentiation were analyzed to see what FGR affected. RA is known to cause expression of FGR which is incorporated into and activates a putative novel cytosolic macromolecular signaling machine(signalsome) that propels differentiation. RA enhances the abundance of signalsome constituents, their associations, and their phosphorylation. The signalsome contains connected nodes that appear as a spine to which the other components are connected. The apparent “nodes” are RAF, LYN, FGR, SLP-76 and CBL. All of these become enriched in the nucleus after RA-treatment. NUMB and VAV appear to provide further scaffolding functions enhanced by RA. RAF in the nucleus complexes with a RARE (retinoic acid-response-element) in the promoter of the blr1 gene, which encodes a serpentine G-protein-coupled-receptor. blr1 transcriptional activation by RA depends on RAF binding. BLR1 expression is necessary to propel RA-induced differentiation, although by itself is not sufficient to cause phenotypic differentiation. Analyzing this signaling process revealed that expression of FGR mimics RA-induced enhancement of the signalsome nodes, enhancing expression of RAF and its phosphorylation, and causing BLR1 expression. Interestingly, for cd38 and blr1, FGR apparently causes expression of genes targeted by RAR/RXR even without RA. FGR thus appears to cause signaling events and phenotypic shift characteristic of RA. In sum, the data indicate that FGR is the “trigger” for RA-induced differentiation. Given the historical perception of FGR as a pro-proliferation, transforming-viral-oncogene, this is a surprising paradigm shift.

## INTRODUCTION

Retinoic acid (RA) is the active metabolite of vitamin A and is a steroid that is necessary for proper growth and development in chordates. During embryonic development, it is a morphogen specifying position along the anterior posterior axis through regulating Hox genes [1]. Its canonical mechanism of action is as a ligand for the ligand-activated transcription factor, RAR, which with its RXR hetero-dimerization partner, binds Retinoic Acid Response Elements (RAREs) in the promoters of targeted genes, thereby releasing co-repressors and attracting co-activators to initiate transcription [2]. In adults, it regulates a variety of tissue and organ functions, including hematopoiesis, immune function, neural function and spermatogenesis, amongst other things. It is ergo of widespread and fundamental biological significance as a regulator of cell proliferation and differentiation. As such, RA has been exploited as a therapeutic modality in the differentiation induction therapy of Acute Promyelocytic Leukemia (APL, French-American British classification FAB M3) where retinoic acid based therapy has caused remission rates of almost 90%, making its mechanism of action and applicability to the treatment of other tumors of great interest in cancer chemotherapy [3]. A major hurdle to molecular insight into its mechanism of action is that RA induces a plethora of changes in gene expression thereby making discerning the events seminal to control of proliferation and differentiation a challenge. It is not known what RA-induced triggering event(s) causes the cascade culminating in cell differentiation [4].

The mechanism of action of RA has been studied in numerous contexts. One of the most prominent experimental venues is the HL-60 human leukemia cell [5, 6]. Derived from a patient with Acute Myeloblastic Leukemia (AML French-American-British classification FAB M2), it is a hematopoietic precursor cell in the myelo-monocytic series that gives rise to either myeloid or monocytic cells. It is thus lineage uncommitted as a lineage bi-potent GM precursor [7–9]. The cells proliferate avidly in culture with a division time of about 24 hrs., making them attractive as an experimental model susceptible to genetic engineering where ample material for molecular analysis can be generated. Over about 2 division cycles, RA causes the cells to undergo myeloid differentiation, whereas 1,25-dihydroxy vitamin D3 causes monocytic differentiation. RA-induced differentiation presents as a progression that is betrayed by sequential expression of cell surface and functional markers, CD38, CD11b, ROS (reactive oxygen species) production which is inducible, and G1/0 cell cycle arrest, culminating in the appearance of mature myeloid cells [10, 11] . The process is driven by MAPK pathway signaling reflecting activation of a RAF/MEK/ERK axis imbedded in a cytosolic signaling machine, a putative signalsome [12–14]. RA activates the signalsome by inducing expression of FGR which is incorporated into the signalsome to activate it. Activation is associated with enhanced expression of its constituents and their phosphorylation [15]. The signalsome includes an ensemble of signaling molecules typically associated with MAPK pathway signaling, including RAF, LYN, FGR, SLP-76 and CBL, which form a string of connected nodes where other components connect as detected by immunoprecipitation and FRET [15–19]. RAF, LYN, FGR, SLP-76 and CBL become enriched in the nucleus, ostensibly reflecting nuclear translocation due to signalsome activation [20–22]. RAF in the nucleus binds NFATc3 in the promotor of the blr1 gene, which encodes a membrane serpentine receptor aka CXCR5, and this is associated with transcriptional activation [22]. BLR1 expression occurs early and is necessary to propel RA-induced differentiation since its KO eliminates RA-induced expression of all aforementioned differentiation markers [23, 24]. This scheme, which is a significant augmentation of the historically dominant paradigm for mechanism of action of RA, occurs amidst a myriad of other RA-induced cellular protein changes, any of which might potentially influence it or even supplant it. However, FGR appears to play a central role as posited in this scheme [15]. Interestingly, chronic exposure of these cells to RA results in resistance to RA, reminiscent of the clinical resistance to RA occurring when APL patients relapse after initially successful RA-treatment, where FGR now fails to be induced by RA [25, 26].

FGR is a member of the Src family of protein tyrosine kinases. It is historically perceived as a viral oncogene with pro-proliferation effects via positive effects on cellular mitogenic signaling, hence it is considered a transforming onco-protein [27]. It is a plasma membrane protein [28]. It notably acts downstream of receptors to facilitate signaling [28–33]. In addition to transformation, it also functions, acting as a positive or negative regulator, in various normal contexts including migration, adhesion, and signal transduction, such as from the membrane to the actin cytoskeleton. It has SH2 and SH3 domains to enable its interactions with the phospho-tyrosine-containing motifs or proline-rich motifs, respectively, of partnering proteins [34]. It has ergo, widespread deployment as an intermediary for regulating cellular signaling [35–37]. Of note and relevance to the present study, it has been found to cause phosphorylation of CBL and VAV [15, 38–40].

The results of the present study augment paradigms on the mechanism of action of RA and FGR. RA is a well-known inducer of cell differentiation, and we now find that ectopic expression by stable transfection of FGR in HL-60 human myeloblastic leukemia GM-precursor cells by itself resulted in a phenotypic shift that was characteristic of differentiation induced by RA. In particular, expression of FGR caused expression of the cell surface differentiation markers, CD38 and CD11b, the functional differentiation marker, ROS, and the cell cycle regulator, p27(kip1). FGR thus supplanted RA in inducing cell differentiation. This appeared to reflect the effect of FGR on signaling that propels RA-induced differentiation. Expression of FGR enhanced the expression of a suite of proteins in a putative signalsome that promotes cell differentiation, mimicking the effects of RA. These include RAF, LYN, SLP-76 and CBL, all of which have been previously implicated in promoting RA-induced differentiation. FGR is known to bind and activate this signalsome, suggesting that RA-induced FGR expression is a seminal event for causing cell differentiation. Expression of FGR also mimicked RA in causing enhancement of VAV, a GEF and adaptor thought necessary for myelopoiesis, and NUMB, an adaptor that regulates binary cell fate decisions during embryonic development; both of which appear to function as scaffolds in the signalsome, depending on their phosphorylation. Also analogous to RA, FGR expression caused expression of BLR1 (aka CXCR5), a serpentine cell surface/membrane receptor, expression of which is known to be necessary for RA-induced differentiation. It is intriguing that CD38 and BLR1 are both genes known to be positively regulated by RAREs, but expression was turned on without RA by FGR. Interestingly, treating the FGR stable transfectants with RA did not appreciably further increase the expression of these key signalsome components or the expression of differentiation markers, consistent with the notion that FGR had effectively usurped/subsumed the function of RA. Among the many proteins that have their expression affected by RA in the process of cell differentiation, FGR thus appears to function as the trigger that causes the signaling and consequential phenotypic shift characteristic of RA-induced differentiation. It is striking that a single molecule can do this, the more so that it is historically a transforming onco-protein.

## MATERIALS AND METHODS

### Cell culture and treatments

Human myeloblastic leukemia cells (HL-60) were grown in a humidified environment of 5% CO2 at 37°C and maintained in RPMI 1640 (Invitrogen, Carlsbad, CA) supplemented by 1% antibiotic/antimycotic (Sigma, St. Louis, MO, USA) and 5% heat-inactivated bovine fetal serum (Hyclone, Logan, UT, USA). At a density of 0.1 × 10^6^ cells/mL, experimental cultures were initiated for lysate collection 72 hours after treatment for all experiments. The hemocytometer and 0.2% trypan blue (Invitrogen, Carlsbad, CA, USA) are used for the dye exclusion assay of cell growth and viability. The same RA (Sigma) treatment conditions (0 μM or 1 μM) were used for all cells and lysate obtained. At least three biological replicates of each experiment were performed. pWZL-Neo-Myr-Flag-DEST was a gift from Jean Zhao (Addgene plasmid # 15300; http://n2t.net/addgene:15300; RRID: Addgene_15300) [41]. After plasmid isolation 2 x10^12^ plasmid copies and 4x10^6^ cells in 0.4 ml of RPMI 1640 without serum were mixed together. Electroporation was done using (Gene Pulser; Bio-Rad Laboratories, Hercules, CA) 300 V and 500 F capacitance in a 0.4-cm electrode gap cuvette. After electroporation, the cells were cultured in a serum-supplemented medium for 2 days to allow their recovery from electroporation. All cells were then harvested and resuspended in a fresh serum-supplemented medium containing 1 mg/ml active G418. The total pool of cells derived from the electroporation was thus subject to selection. Pooled transfectants were used to obviate clonal variation. By 25 days, the transfected cells resumed growth, but at a slower rate compared with the wild-type HL-60. After 3 weeks of selection cultures will be amplified for one week and then used for experiments. After thawing cryogenic-preserved stocks, cells were subject to selection for three weeks and then amplified for one week prior to experimental use. The cell line Human myeloblastic leukemia cells (HL-60) was derived from the original isolates, a generous gift of Dr. Robert Gallagher and maintained in this laboratory, certified and tested for mycotoxin by Biosynthesis, Lewisville, TX, USA, in August 2017.

### Antibodies and reagents

Antibodies for flow cytometric analysis, PE-conjugated CD38 (clone HIT2) and APC-conjugated CD11b (clone ICRF44) conjugated with allophycocyanin (APC), were from BD Biosciences (San Jose, CA, USA). SLP-76, Lyn, Fgr, Vav1, p-tyr, HRP anti-mouse and anti-rabbit antibodies were purchased from Cell Signaling Technologies (Danvers, MA, USA). Anti-c-Cbl (clone C-15, catalogue number sc-170, lot H0414) was purchased from Santa Cruz Biotechnology (Santa Cruz, CA, USA). NUMB Antibody catalogue number (703078) was from Thermo Fisher Scientific (Waltham, MA, USA). The c-Raf antibody was from BD Biosciences (San Jose, CA, USA). Protease and phosphatase inhibitors were purchased from Sigma (St. Louis, MO, USA). Protein G magnetic beads used for immunoprecipitation were from Millipore (Billerica, MA, USA).

### CD38, CD11b quantification and phenotypic analysis

Immunostaining for CD38 and CD11b was performed as previously stated [17] and fluorescence was detected using a Becton Dickinson LSR II flow cytometer (San Jose, CA, USA). Flow cytometric phenotypic analysis gating for positives was set to exclude 95% of the untreated cells in HL-60 wildtype and FGR O.E samples.

### Measurement of respiratory burst (inducible oxidative metabolism)

0.5 × 10^6^ cells from HL-60 were harvested by centrifugation and resuspended in PBS 200 μL with 10 μM 5-(and-6)-chloromethyl-2′,7′—containing Acetyl ester of dichlorodihydrofluorescein diacetate (H2 -DCF, Eugene, Molecular Probes, OR) and 0.4 μg/mL 12-O-tetradecanoylphorbol-13-acetate (TPA, Sigma). samples were incubated at 37°C in a humidified atmosphere of 5% CO_2_ for 20 minutes. Flow cytometric analysis was performed (BD LSRII flow cytometer) using 488-nm laser excitation with emission collected through a 505 long-pass filter. The fluorescence Intensity shift in response to TPA was used to evaluate the percent of cells with inducible generation of superoxide. Gates to assess the percentage of positive cells were set to exclude 95 percent of the control cells that did not get TPA to ca respiratory burst distinguishing mature cells. Samples with or without TPA of cells that have not been RA-treated and without TPA of RA-treated cells showed indistinguishable DCF fluorescence histograms [10].

### Cell-cycle quantification

A total of 1 × 10^6^ cells were centrifuged and resuspended in 200 μL of cold propidium iodide (PI) hypotonic staining solution containing 50 μg/mL of propidium iodine, 1 μL/mL of triton X-100 and 1 mg/mL of sodium citrate. Cells were incubated for 1 hour at 4°C and analyzed by flow cytometry using 488-nm excitation and emission measured with a 576/26 band-pass filter (BD LSRII). Doublets were classified and removed from the study by a PI signal width versus area map [42].

### Western blotting

Cell fractionation was performed with the NE-PER kit (Pierce) in accordance with the manufacturer’s instructions, with the addition of protease and phosphatase inhibitors, and all lysates were stored at −80°C before use. After lysate collection, cell debris was cleared by centrifugation at 13,000 rpm for at least 10 minutes, and protein concentrations were quantified using the BCA (Pierce) assay.

For Western blotting, 25 μg protein per lane was resolved on a 12 percent polyacrylamide gel. The electro-transfer was done at 400 mA for 1 hour. The membranes were blocked in milk for 1 hour before adding the primary antibody and was incubated overnight at 4°C. The images were captured on a ChemiDoc XRS Bio-Rad Molecular Imager and analyzed using the Image J program.

Densitometric values were measured for each Western Blot band. The values were then normalized using Image J to the loading control for the lane. In the bar graphs, the lowest normalized value is arbitrarily set to one and the values for the other bands were normalized to it and thus compared to the lowest value, which is typically the same as the untreated control, unless the signal was not detectable in which case the lowest detectable signal was used. Values from at least three biological repeats were computed using GraphPad Prism 6.01 and statistically evaluated.

### Statistical analysis

The experiments were triplicate biological replicates, and the findings are shown as mean and standard deviation (SD). For the estimation of the difference between two classes, a two-tailed paired *t* test was used. A *p* value less than 0.05 was considered significant.

## RESULTS

### FGR EXPRESSION CAUSED CELL DIFFERENTIATION

Retinoic acid is a major regulator of embryonic development, a process of cellular decisions to choose phenotypes and proliferate or arrest that involves a complex interrelated set of decision points. HL-60 cells are lineage uncommitted cells that make only two decisions, namely, to grow or not and which of two lineages to differentiate along; in that sense they might be thought of as an abstraction of broader and more complex RA-guided developmental biology. RA causes these cells to elect the myeloid lineage and undergo G1 growth retardation, betrayed by expression of differentiation markers that are well known since hematological cell markers have been historically so well established. The major ones for myeloid lineage differentiation are CD38, CD11b, ROS, and for growth arrest, p27(kip1).

#### Stable transfectants achieve FGR expression comparable to level induced by RA

Stable transfectants ectopically expressing FGR were created. Wild type (wt) HL-60 cells were stably transfected to express FGR using electroporation of a retroviral expression vector (pWZL Neo Myr Flag FGR) with a dominant selectable marker (Neo) and G418 selection. Transfectants were pooled to avoid clonal bias. The wild type of parental cells and the FGR stable transfectants, untreated and RA-treated (1.0 microM for 72 hrs), were subject to Western analysis probing for FGR (FIGURE 1). The Western blot shows that there is no expression in untreated wild type cells, and there is pronounced expression in the stable transfectants. The extent of expression was comparable to that typical of levels induced by RA in wild type cells. RA (1 microM) did not cause any gross enhancement of FGR in the stable transfectants. These stable FGR transfectants were used in the subsequent experiments that analyzed their phenotype and signaling attributes.

**Figure 1:**
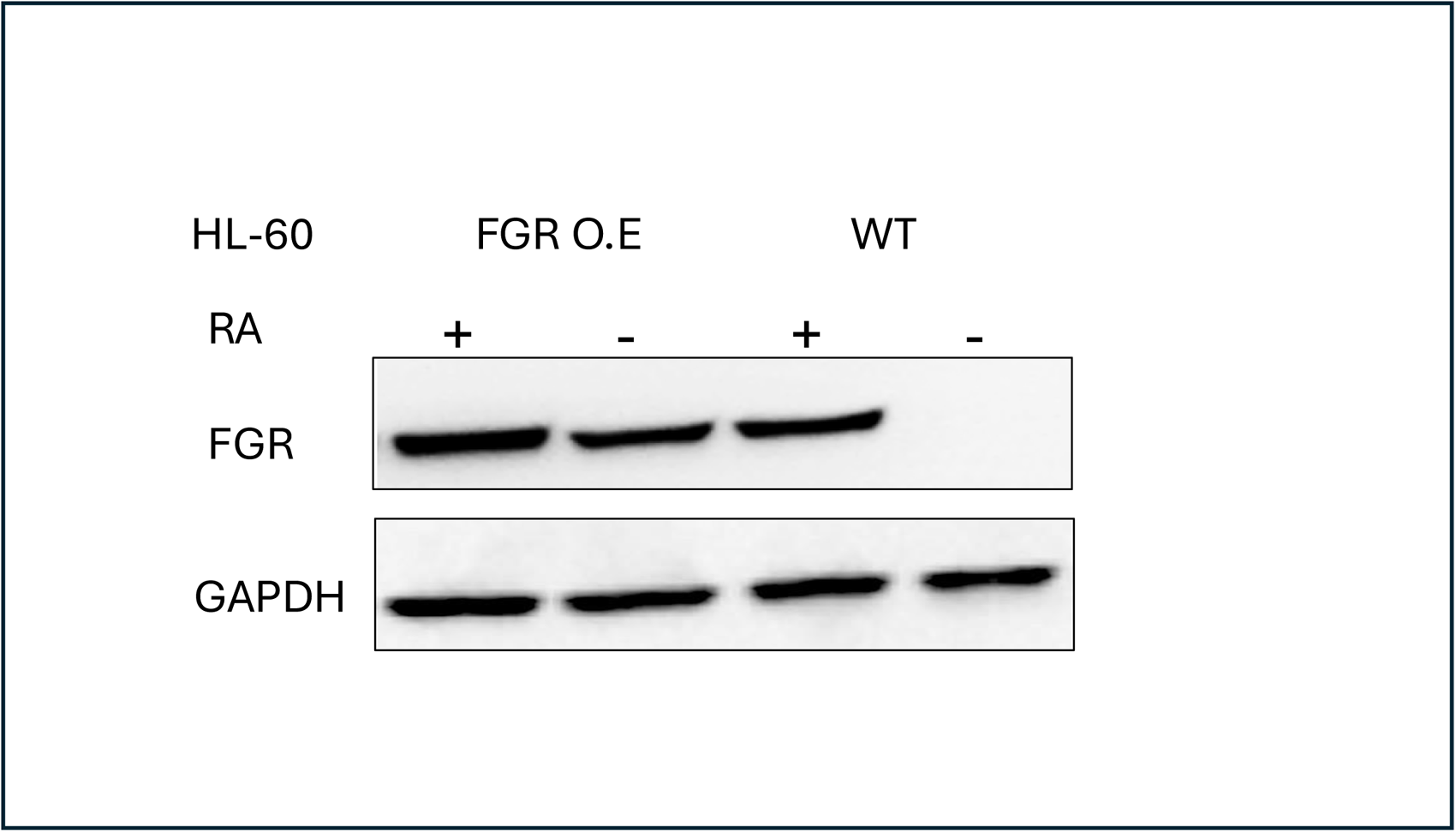
FGR Western blot analysis of HL-60 wt and FGR O.E cells untreated and treated with RA. Wild-type and FGR O.E HL-60 cells were untreated (control) or treated for 72 h with RA (1 μM) as indicated. 25 μg of lysate per lane was resolved by SDS PAGE and electro-transferred to membranes. Membrane images for each protein are cropped to show only the band of interest. For all Western blots, densitometric analysis of 3 or more biological repeats were quantified using ImageJ and shown in Figure S10.

#### FGR stable transfectants express cell surface differentiation marker CD38

Unlike wt parental cells, FGR stable transfectants express CD38. CD38 is a cell surface marker for RA-induced myeloid differentiation. It is an ecto enzyme receptor for NAD metabolism that also originates MAPK pathway signaling. In the RA-induced progression of wt HL-60 cells from myeloblastic to mature myeloid cells, CD38 is an early marker that is apparent in most cells by about 12 hrs of treatment. Cell membrane CD38 expression was analyzed using a phycoerythrin conjugated antibody against CD38 that was detected by flow cytometry. The flow cytometric histogram shows relative number of cells versus their expression level for wt cells and stable transfectants that were untreated or RA-treated (1.0 microM for 72 hrs) (FIGURE 2). At least 10,000 cells were analyzed for each case. Wt HL-60 were negative for expression, whereas the FGR stable transfectants were positive. The CD38 expression level achieved in the stable transfectants was comparable to that of RA-treated wild type cells. Treating the FGR transfectants with RA did not cause any significant further increase in CD38 expression level.

**Figure 2:**
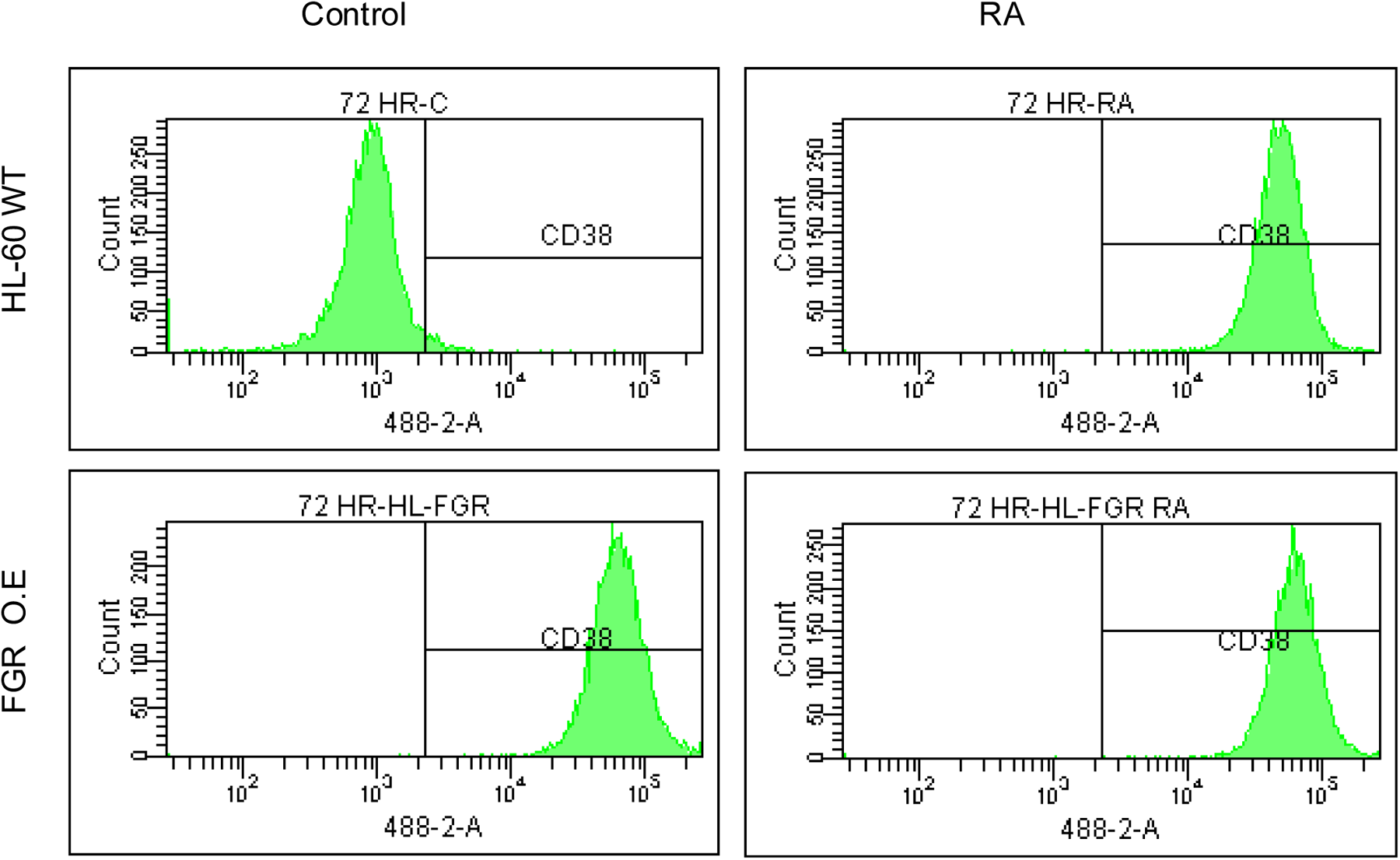
Phenotypic cell surface differentiation marker analysis of HL-60 wt and FGR O.E cells untreated and treated with RA. HL-60 cells were cultured in the absence (control) or presence of 1 μM RA as indicated. CD38 expression was assessed by flow cytometry following 72 h treatment period. Gating to discriminate positive cells was set to exclude 95% of untreated controls. Quantification of 3 or more biological repeats is in Figure S19.

#### FGR stable transfectants express CD11b cell surface differentiation marker

Unlike wt parental cells, FGR stable transfectants express CD11b. CD11b is a cell surface marker for RA-induced myeloid differentiation. It is an integrin receptor, and a complement receptor, and can originate MAPK pathway signaling. In the RA-induced progression of cell differentiation, it is a marker of later progression that follows CD38 expression. For wt cells and stable transfectants that were untreated, or RA treated (1.0 microM for 72 hrs), cell surface CD11b expression was analyzed by flow cytometry using a fluorescent FITC conjugated antibody against CD11b (FIGURE 3). The flow cytometric histograms showing relative number of cells versus expression level show that while wt cells are negative, the FGR transfectants are positive. The expression level in the transfectants was comparable to that of wt cells treated with RA. Expression in the transfectants was not significantly enhanced by treatment with RA. Ectopic FGR expression thus resulted in cells displaying two cell surface markers that characterize RA-induced differentiation.

**Figure 3:**
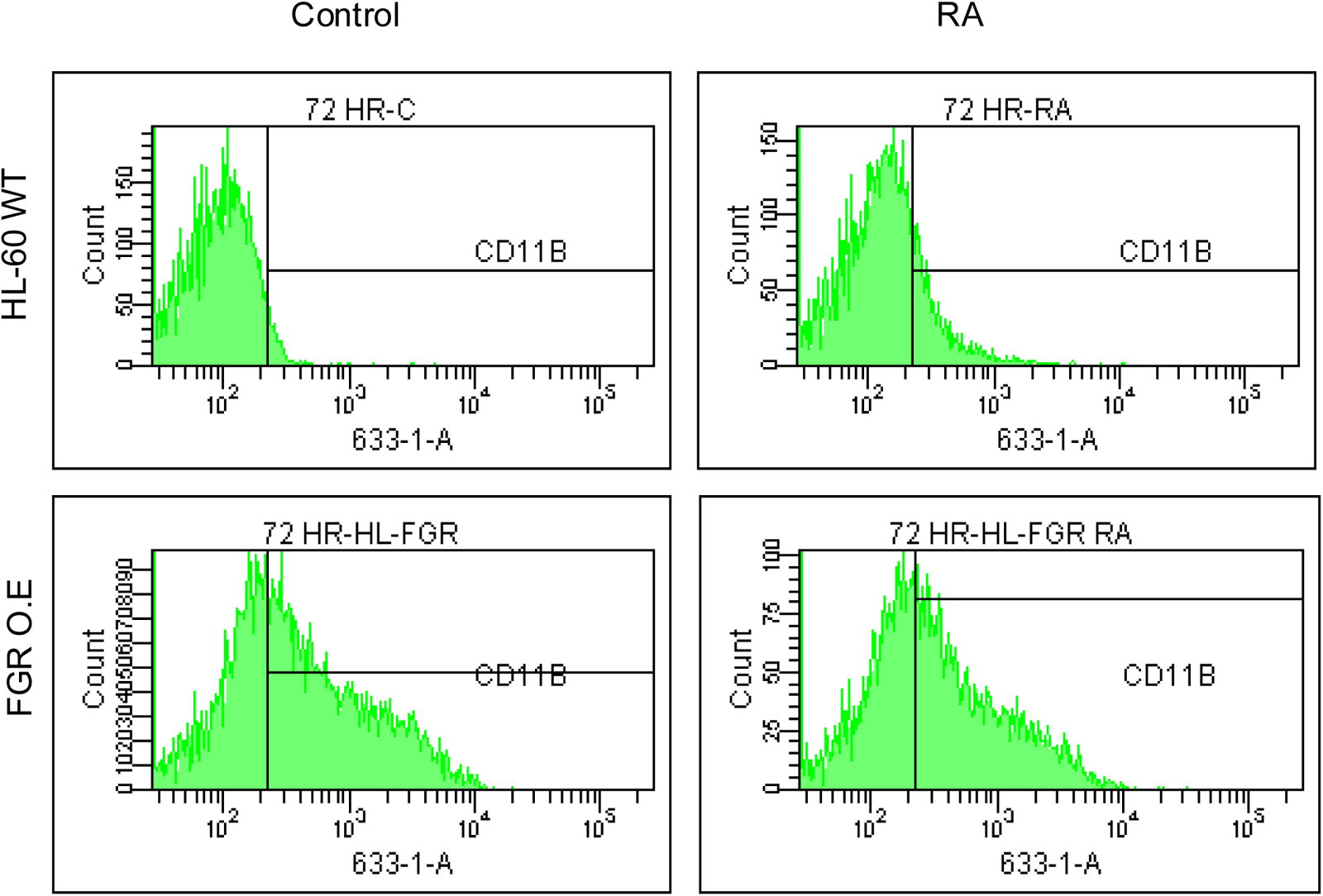
Phenotypic cell surface differentiation marker analysis of HL-60 wt and FGR O.E cells untreated and treated with RA. CD11b expression was assessed by flow cytometry after 72 h treatment periods. Gating to discriminate positive cells was 95% of untreated controls. Quantification of 3 or more biological repeats is in Figure S20.

As expected for myeloid differentiation, expression of the monocyte/macrophage cell surface differentiation marker, CD14, was not detected by flow cytometry in FGR stable transfectants, matching wt parental cells treated with RA which were also negative (data not shown).

#### FGR stable transfectants express ROS

Unlike wt parental cells, FGR stable transfectants produce ROS (reactive oxygen species) that is inducible. Production of reactive oxygen species, superoxide, is a functional differentiation marker of mature myeloid cells. ROS production is inducible, a telltale of the inducible oxidative metabolism that characterizes mature myeloid cells. For wt cells and stable transfectants that were untreated or RA-treated (1.0 microM for 72 hrs), ROS was assayed using 5-(and-6) chloromethyl-2′,7′-dichlorodihydro–fluorescein diacetate acetyl ester (H2-DCF) oxidation detected by flow cytometry (FIGURE 4). The flow cytometric histogram shows the relative number of cells versus fluorescence. TPA was used to cause inducible superoxide production. DMSO was the carrier for the TPA; and DMSO-treated cells were the carrier control for assaying TPA inducible oxidative metabolism. Whereas wt cells did not make detectable ROS, as expected for cells that had not differentiated, the FGR transfectants were producing ROS. TPA failed to induce much more ROS production in wt cells as expected for cells that had not differentiated, but FGR transfectants produced more ROS after TPA treatment. The FGR transfectants thus expressed ROS, and it was TPA-inducible, characteristic of differentiated cells. After RA-treatment, wt cells showed TPA-inducible ROS as expected of differentiated cells. ROS production induced by TPA in FGR transfectants was comparable to that in RA-treated wt cells. The FGR transfectants thus expressed a functional differentiation marker, namely ROS production, characterizing cell differentiation in addition to the two cell surface markers.

**Figure 4:**
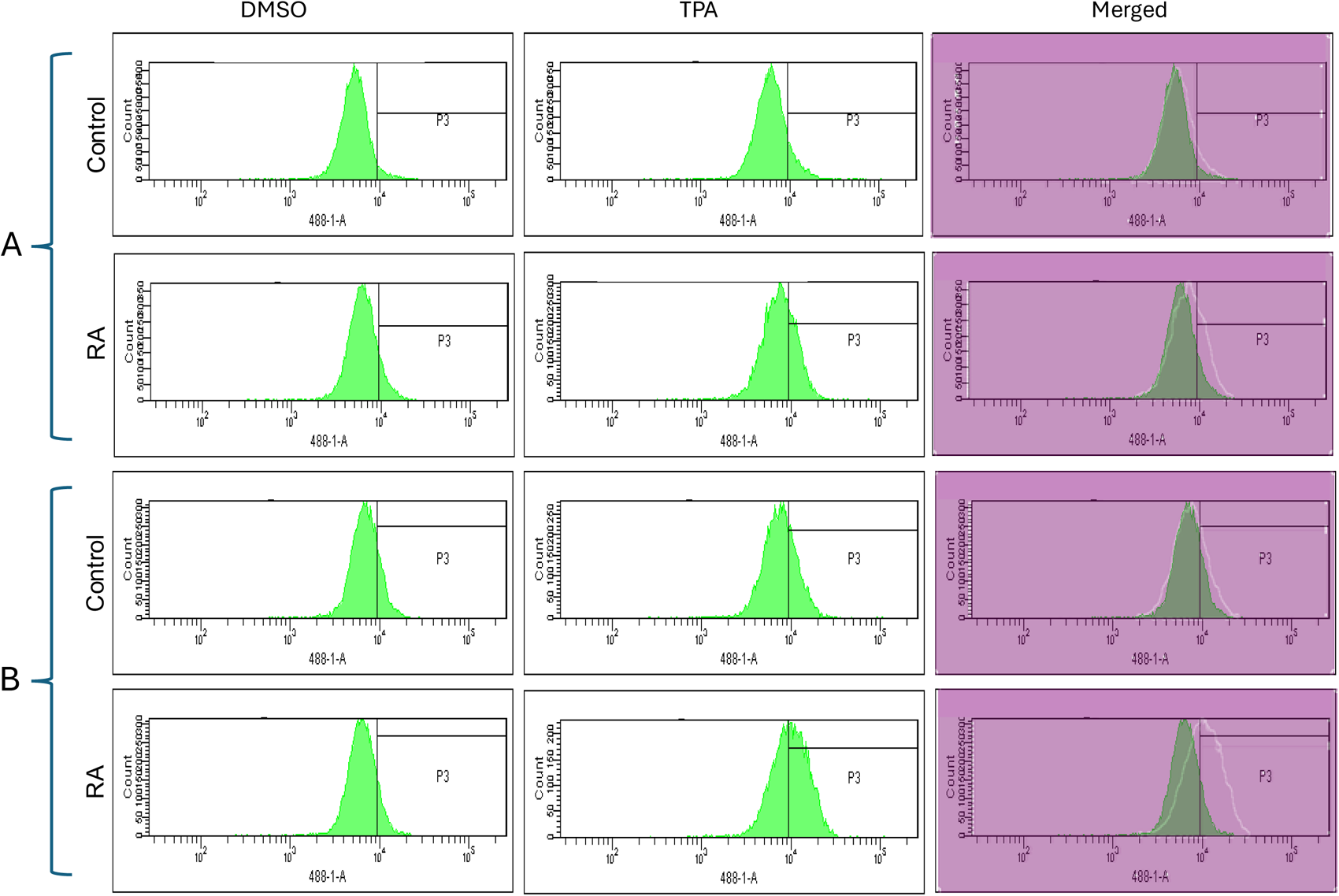
Functional differentiation marker analysis of HL-60 wt and FGR O.E cells untreated and treated with RA measured by TPA-induced respiratory burst. HL-60 WT (parental wildtype) cells were cultured in the absence (control) or presence of 1 μM RA as indicated. FGR O.E cells were cultured in the absence or presence of 1 μM RA as indicated. Respiratory burst was analyzed by measuring reactive oxygen species (ROS) production by flow cytometry using the 2′,7′-dichlorofluorescein (DCF) assay for DMSO carrier control and TPA induced cells. Gates shown in the histograms were set to exclude 95% of the DMSO-treated control population (carrier control) for each culture condition. For each of the 4 cases, WT and FGR that were control and RA-treated, TPA-treated samples show induced ROS. Inducible ROS production is betrayed by the shift in the TPA histogram compared to the DMSO histogram shown in the merged histogram. Quantification of 3 or more biological repeats is in Figure S21.

#### FGR stable transfectants express p27 CDKI

Expression of FGR in stable transfectants results in expression of p27. P27 is a CDKI (Cyclin Dependent Kinase Inhibitor). It inhibits G1 cyclins to retard cell cycle progression in G1. Wt HL-60 cells do not express p27. Western analysis of wt cells and FGR transfectants that were untreated or treated with RA (1.0 microM for 72 hrs) showed that while wt cells were negative, the FGR transfectants were positive (FIGURE 5). The expression level of p27 in the FGR transfectants was comparable to that induced by RA in wt cells. Treating the FGR transfectants with RA did not cause a detectable increase in expression. Hence, the expression of FGR caused expression of a cell cycle regulator characteristic of RA treatment. As expected, population growth of the cells mirrored this. Flow cytometric DNA histograms of cells after hypotonic propidium iodide staining showed that the FGR transfectants were enriched for relative number of G1/0 cells compared to the wt cells (FIGURE 6). In sum, FGR expression in stable transfectants caused the expression of an ensemble of cell surface and functional differentiation markers characteristic of RA-induced differentiation, specifically CD38, CD11b, ROS, and p27. FGR has thus effectively caused cell differentiation akin to RA.

**Figure 5:**
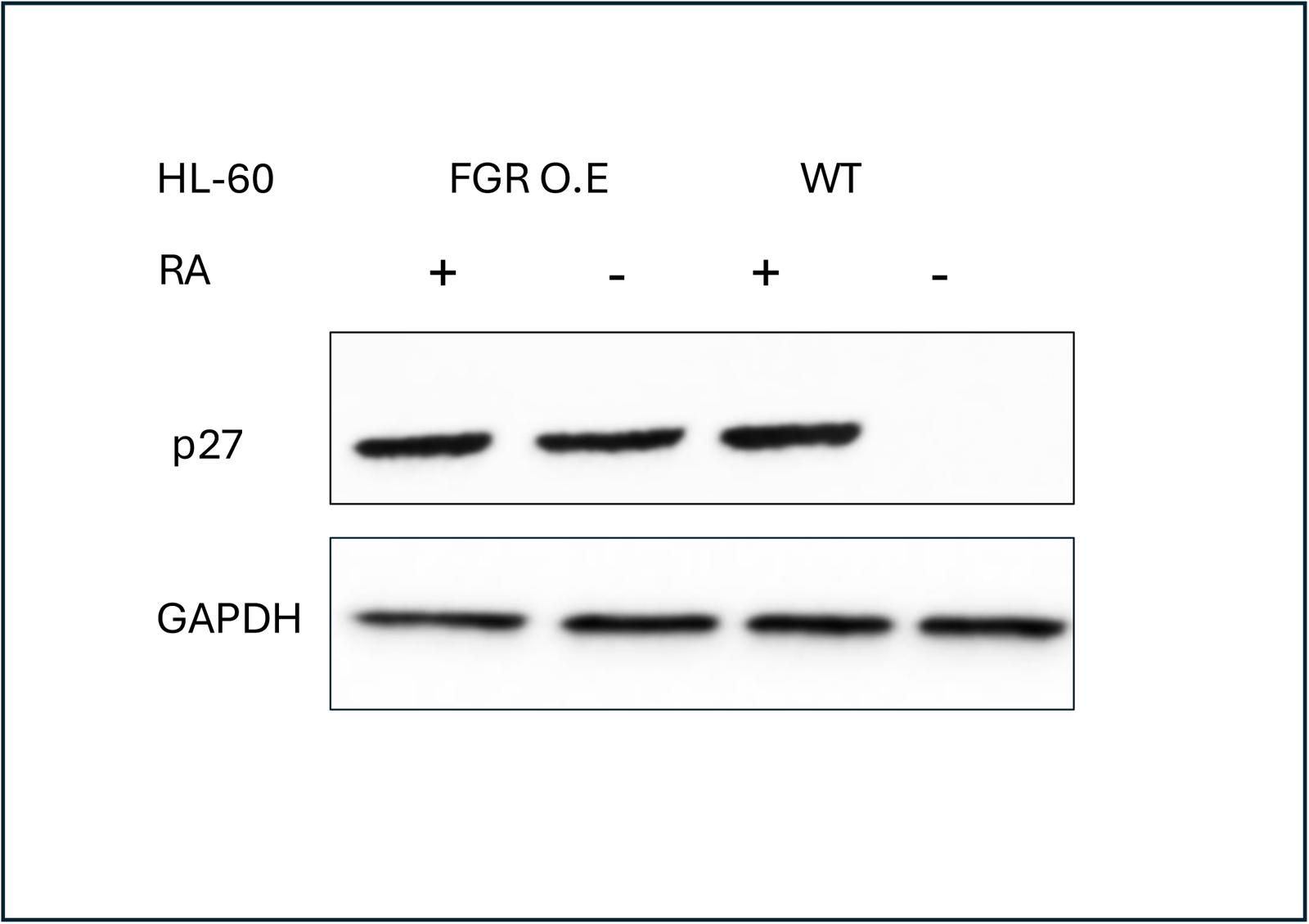
p27 Western blot analysis of HL-60 wt and FGR O.E cells untreated and treated with RA. Wild-type and FGR O.E HL-60 cells were untreated (control) or treated for 72 h with RA (1 μM) as indicated. 25 μg of lysate per lane was resolved by SDS PAGE and electro-transferred to membranes. Membrane images for each protein are cropped to show only the band of interest. For all Western blots, densitometric analysis of 3 or more biological repeats were quantified using ImageJ and shown in Figure S11.

**Figure 6:**
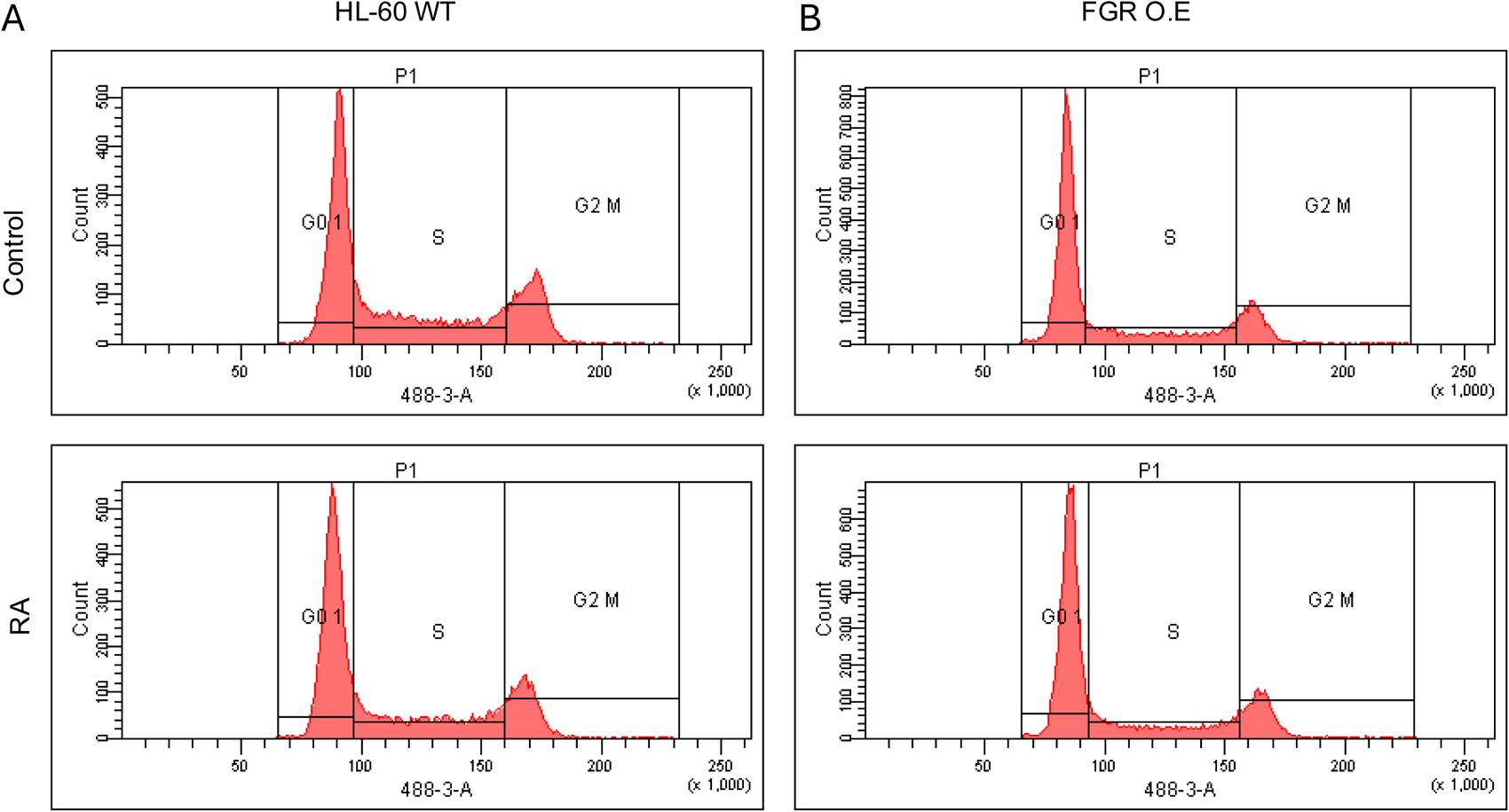
Cell cycle analysis of HL-60 WT and FGR O.E cells. DNA histograms show FGR transfectants were enriched for relative number of G1/0 cells compared to the wt cells. Wild-type and FGR O.E cell lines were cultured for 72 h without (untreated control) or with 1 μM RA as indicated. Cell cycle distribution showing the percentage of cells in G1/G0 was analyzed using flow cytometry with propidium iodide staining at 72 h. Gates define the G1, S, and G2/M subpopulations (left to right). G1/0 arrest is indicated by an increase in the G1 peak for FGR O.E compared to wt cells. Quantification of 3 or more biological repeats is in Figure S22.

A second FGR transfection done as a biological repeat resulted in FGR stable transfectants again expressing these differentiation markers (FIGURE S1). Two biological repeats ergo both showed FGR caused differentiation. Clonal bias in both cases was obviated by using pooled transfected cells.

While it is only anecdotal, it is interesting to note that it appears the second transfectant expressed FGR at a lower level than the first, but unlike the first transfectants RA-treatment obviously enhanced it. And associated with this, expression of the phenotypic markers, CD38, CD11b, ROS and G0, already enhanced by FGR expression, were also further enhanced by RA-treatment (FIGURE S2-S5). This would be consistent with differentiation being driven by FGR expression.

### FGR EXPRESSION ENHANCED SIGNALING MOLECULES THAT PROPEL DIFFERENTIATION

RA-induced differentiation of HL-60 cells is driven by a macromolecular signaling machine, a putative signalsome, in which the RAF/MEK/ERK axis is imbedded. A suite of molecules in the signalsome has been identified as connected nodal points that other signalsome components are attached to. They are up regulated by RA and positively regulate differentiation as revealed by ectopic expression and/or KO. These are RAF, FGR, LYN, SLP-76, and CBL. In the signalsome, VAV and NUMB are implicated as scaffolds that are upregulated by RA. An important function of this signaling is to induce the expression of the BLR1 receptor which is necessary for cells to differentiate.

#### FGR expression enhanced expression of RAF (pS259)

FGR caused increased expression of phosphorylated RAF in transfectants compared to the parental wt cells. In wt cells, RA causes upregulation of RAF and its phosphorylation at several sites, S259 in the CR2 region that contains a recognition site for 143-3, a signalsome component, being a prominent one. RAF activity, embodied in the CR3 region, is known to propel RA-induced differentiation. pS259, which has been reported paradoxically as both a positive and negative regulatory site in other contexts, is used here as a telltale of RA-response. Wt cells and FGR transfectants were untreated or treated with RA (1.0 microM for 72 hrs), and whole cell lysates harvested for Western blotting which was probed for pRAF (pS259) (FIGURE 7). Comparing the FGR transfectants to parental wt cells shows that expression of FGR had caused increased expression of pRAF. The increase in expression met and exceeded that in RA-treated wt cells, where RA caused up-regulation as expected. FGR thus caused up-regulation of pRAF like RA.

**Figure 7:**
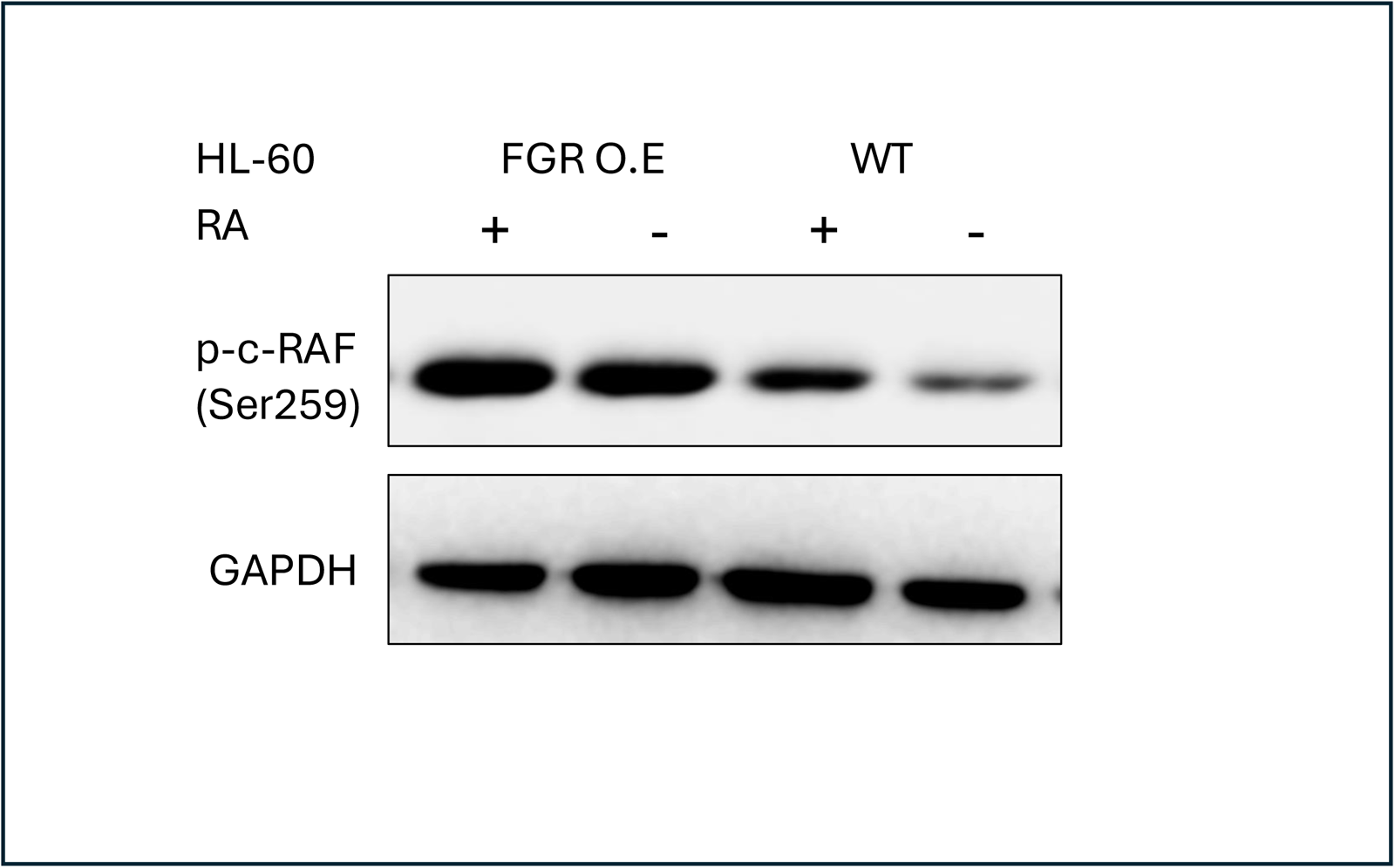
p-c-RAF (Ser259) Western blot analysis of HL-60 wt and FGR O.E cells untreated and treated with RA. Wild-type and Fgr O.E HL-60 cells were untreated (control) or treated for 72 h with RA (1 μM) as indicated. 25 μg of lysate per lane was resolved by SDS PAGE and electro-transferred to membranes. Membrane images for each protein are cropped to show only the band of interest. For all Western blots, densitometric analysis of 3 or more biological repeats were quantified using ImageJ and shown in Figure S12.

#### FGR expression enhanced expression of LYN

FGR caused increased expression of LYN in transfectants compared to wt cells. In wt cells, RA causes up-regulation of LYN, but its regulation of differentiation remains enigmatic. It is associated with both pro-proliferation and pro-differentiation attributes in AML [43]. While a modest KD is inhibitory for RA-induced differentiation signaling, a KO enhances RA-induced signaling and differentiation [16, 44, 45]. Regardless, it is RA-up-regulated, and it regulates RA-induced differentiation – possibly in a dose dependent manner. Wt cells and FGR transfectants were untreated or treated with RA (1.0 microM for 72 hrs), and whole cell lysates harvested for Western blotting, probing for LYN (FIGURE 8). Comparing the FGR transfectants to parental wt cells shows that expression of FGR had caused increased expression of LYN. In wt cells RA caused up-regulation of LYN, as expected. The up-regulated expression in FGR transfectants was comparable to that in RA-treated wt cells. FGR thus caused up-regulation of LYN like RA.

**Figure 8:**
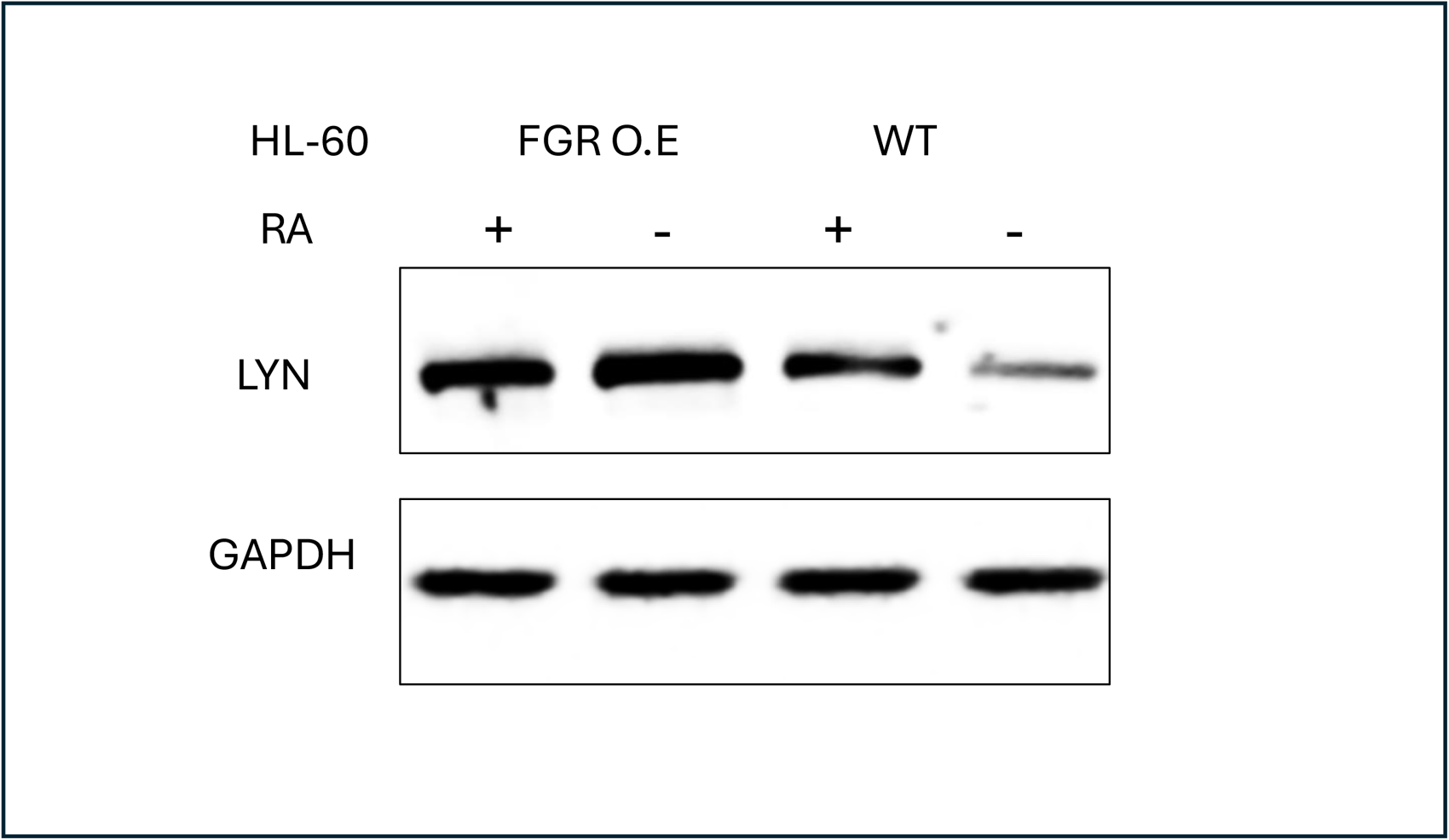
LYN Western blot analysis of HL-60 wt and FGR O.E cells untreated and treated with RA. Wild-type and Fgr O.E HL-60 cells were untreated (control) or treated for 72 h with RA (1 μM) as indicated. 25 μg of lysate per lane was resolved by SDS PAGE and electro-transferred to membranes. Membrane images for each protein are cropped to show only the band of interest. For all Western blots, densitometric analysis of 3 or more biological repeats were quantified using ImageJ and shown in Figure S13.

#### FGR expression enhanced expression of SLP-76

FGR caused increased expression of SLP-76 in transfectants compared to wt cells. In wt cells, RA causes up-regulation of SLP-76, seen as the most prominent RA-induced tyrosine phosphorylation in a p-tyr blot, that was crippled when differentiation was inhibited by FGR KO, implicating it in driving RA-induced differentiation. Wt cells and FGR transfectants were untreated or treated with RA (1.0 microM for 72 hrs), and whole cell lysates harvested for Western blotting, probing for SLP-76 (FIGURE 9). Comparing the FGR transfectants to parental wt cells shows that expression of FGR had caused increased expression of SLP-76. The increased expression was comparable to that in RA-treated wt cells, where RA increased expression as expected. FGR thus caused up-regulation of SLP-76 like RA.

**Figure 9:**
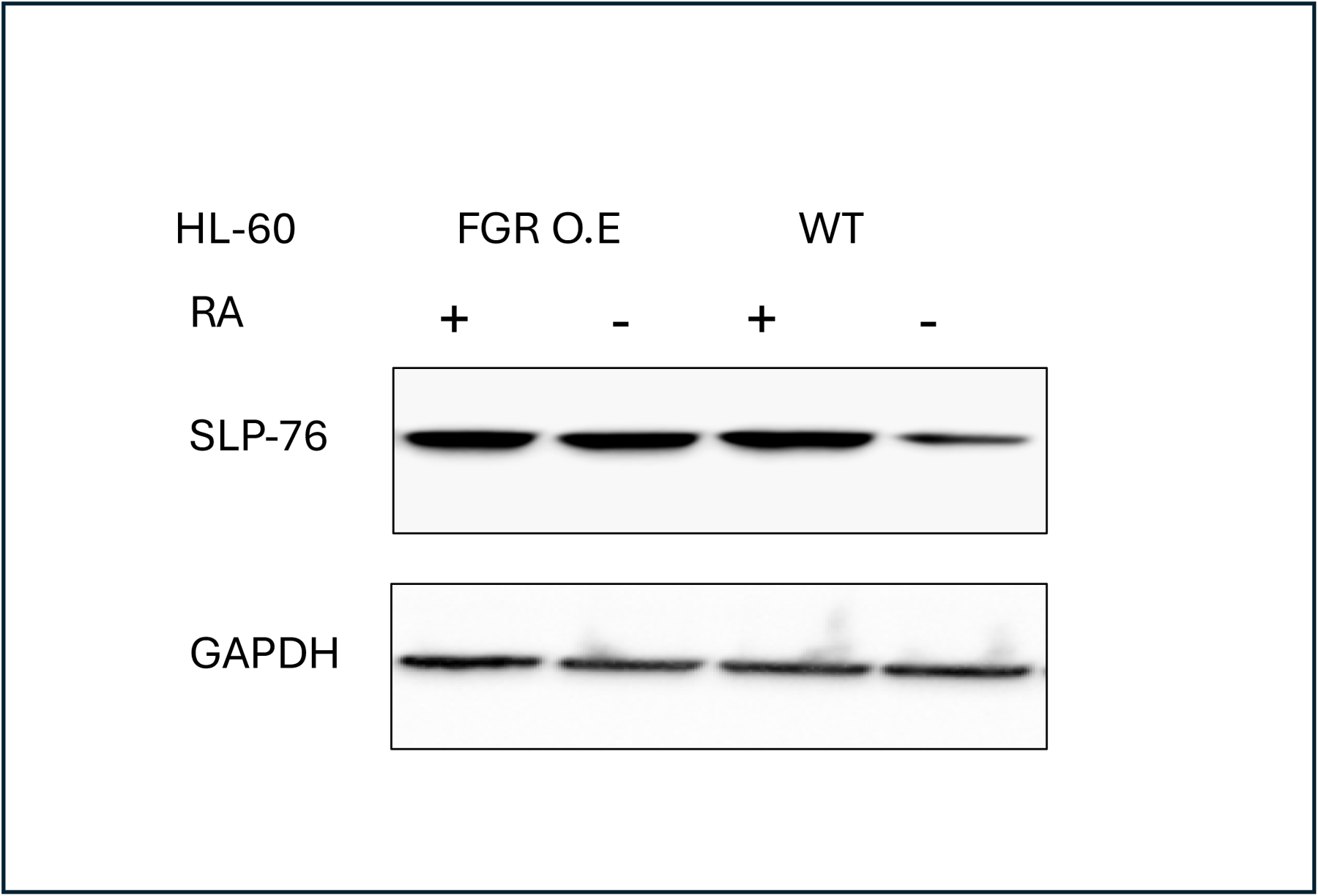
SLP-76 Western blot analysis of HL-60 wt and FGR O.E cells untreated and treated with RA. Wild-type and Fgr O.E HL-60 cells were untreated (control) or treated for 72 h with RA (1 μM) as indicated. 25 μg of lysate per lane was resolved by SDS PAGE and electro-transferred to membranes. Membrane images for each protein are cropped to show only the band of interest. For all Western blots, densitometric analysis of 3 or more biological repeats were quantified using ImageJ and shown in Figure S14.

#### FGR expression enhanced expression of CBL

FGR caused increased expression of CBL compared to wt cells. In wt cells, RA causes a modest enhancement of CBL expression, and ectopically expressed mutant CBL fail to drive differentiation in contrast to wt CBL, implicating it as a regulator of RA-induced differentiation. Wt cells and FGR transfectants were untreated or treated with RA (1.0 microM for 72 hrs), and whole cell lysates harvested for Western blotting, probing for CBL (FIGURE 10). Comparing the FGR transfectants to parental wt cells shows that expression of FGR had caused increased expression of CBL. CBL expression in the transfectants was comparable to that in RA-treated wt cells. FGR transfectants thus had increased CBL comparable to that in RA-treated wt cells.

**Figure 10:**
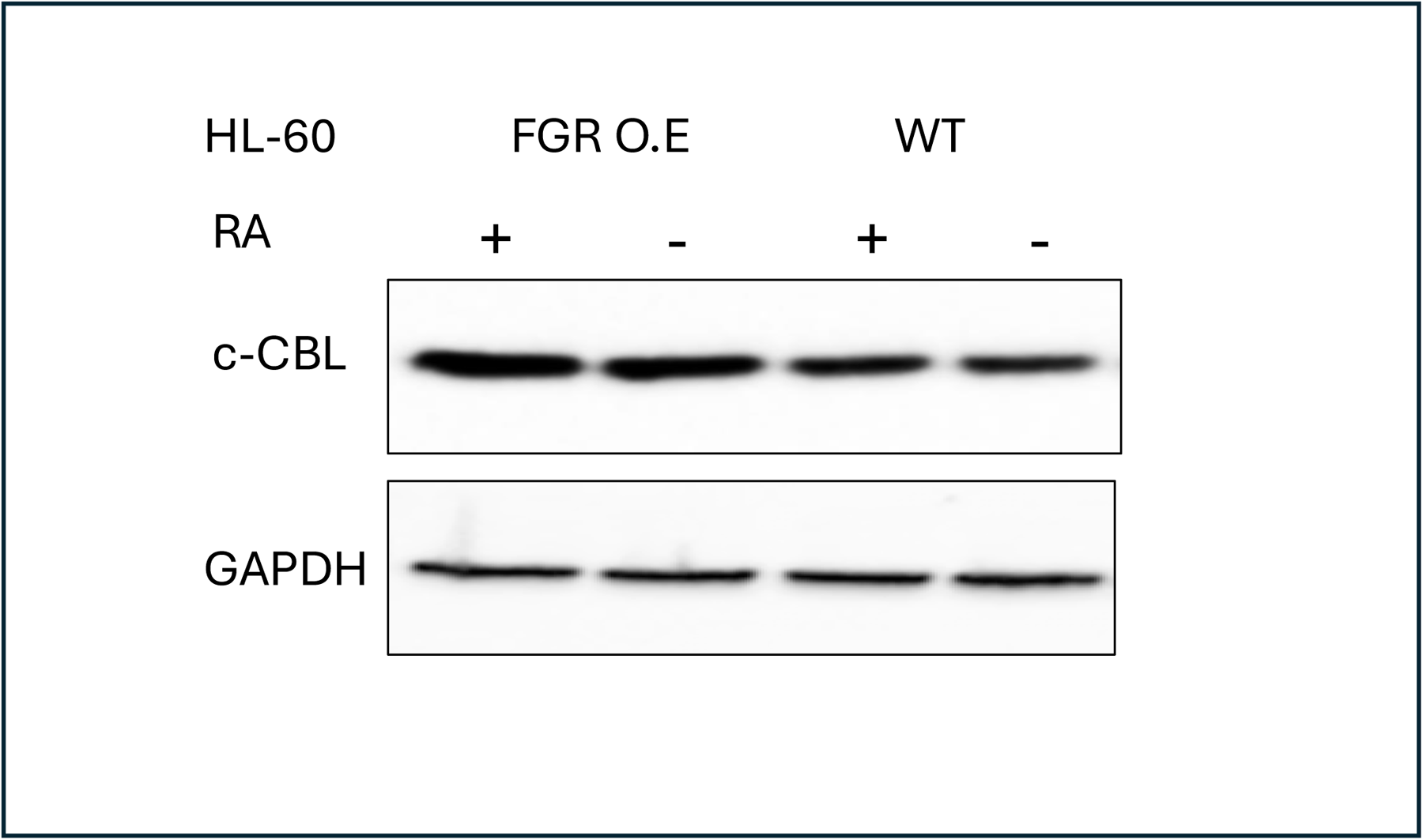
c-CBL Western blot analysis of HL-60 wt and FGR O.E cells untreated and treated with RA. Wild-type and Fgr O.E HL-60 cells were untreated (control) or treated for 72 h with RA (1 μM) as indicated. 25 μg of lysate per lane was resolved by SDS PAGE and electro-transferred to membranes. Membrane images for each protein are cropped to show only the band of interest. For all Western blots, densitometric analysis of 3 or more biological repeats were quantified using ImageJ and shown in Figure S15.

#### FGR expression enhanced expression of VAV

FGR caused enhanced VAV expression compared to wt cells. In wt cells, RA enhances expression of VAV which is a binding partner of CBL, implicating it in regulating RA-induced differentiation. Wt cells and FGR transfectants were untreated or treated with RA (1.0 microM for 72 hrs), and whole cell lysates harvested for Western blotting, probing for VAV (FIGURE 11). FGR transfectants expressed more VAV than wt cells. Expression of VAV in transfectants was comparable to RA-treated wt cells. FGR thus caused increased expression of VAV like RA.

**Figure 11:**
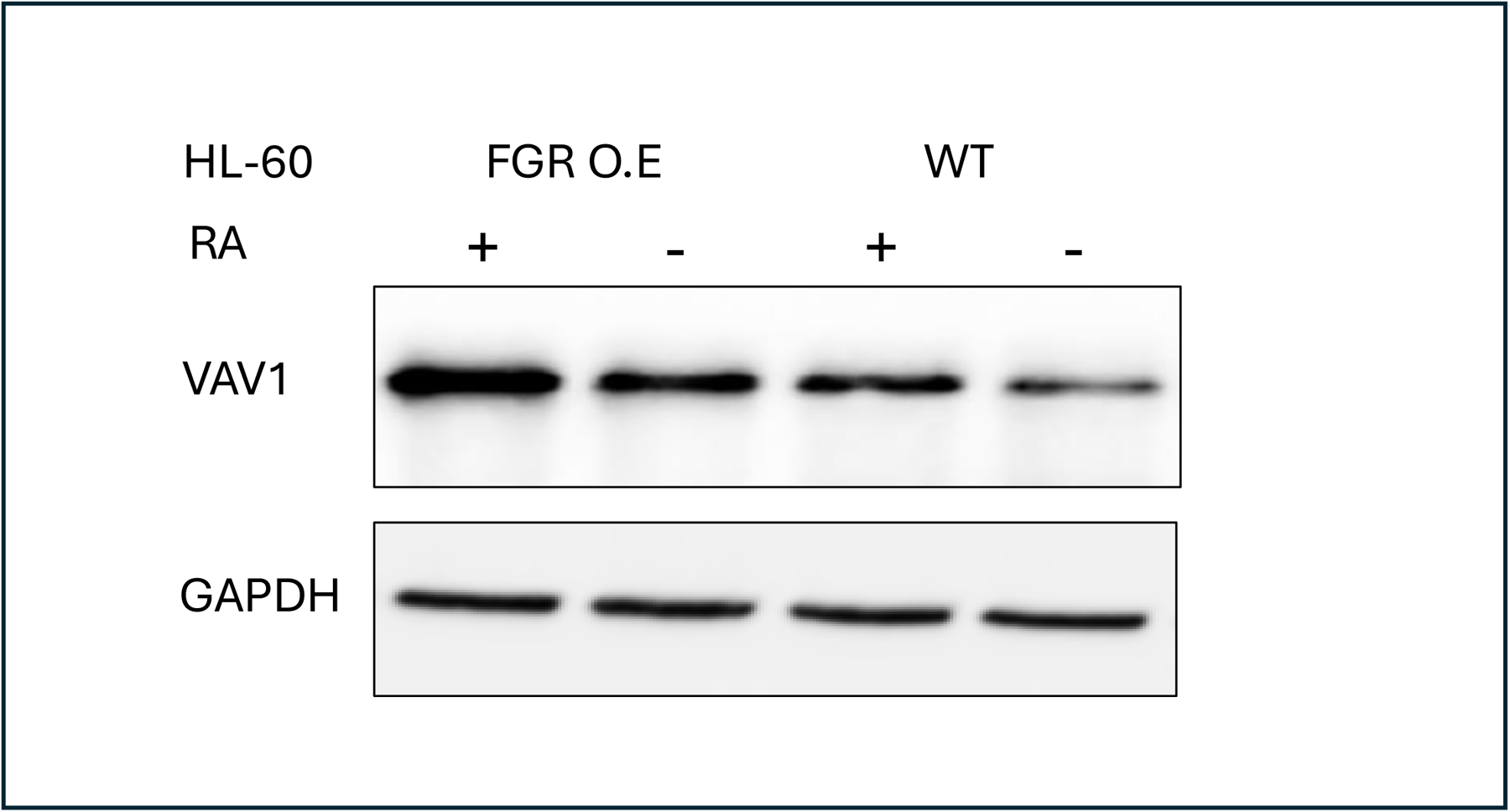
VAV1 Western blot analysis of HL-60 wt and FGR O.E cells untreated and treated with RA. Wild-type and Fgr O.E HL-60 cells were untreated (control) or treated for 72 h with RA (1 μM) as indicated. 25 μg of lysate per lane was resolved by SDS PAGE and electro-transferred to membranes. Membrane images for each protein are cropped to show only the band of interest. For all Western blots, densitometric analysis of 3 or more biological repeats were quantified using ImageJ and shown in Figure S16.

#### FGR expression enhanced expression of NUMB

FGR caused increased expression of NUMB in transfectants compared to expression in wt cells. In wt cells, RA causes increased NUMB expression with attendant FGR dependent phosphorylation, implicating it as a regulator of RA-induced differentiation. NUMB is Ser and Tyr phosphorylated at numerous sites, and the evolutionarily conserved pS276 has been implicated as a regulator of binding to 14-3-3, a component of the signalsome, and asymmetric division. It is used here as a telltale of RA-response. Wt cells and FGR transfectants were untreated or treated with RA (1.0 microM for 72 hrs), and whole cell lysates harvested for Western blotting, probing for pNUMB (pS276) (FIGURE 12). FGR transfectants expressed more pNUMB than wt cells. The expression in FGR transfectants was comparable to RA-treated wt cells. Expression of FGR thus caused increased pNUMB expression like RA.

**Figure 12:**
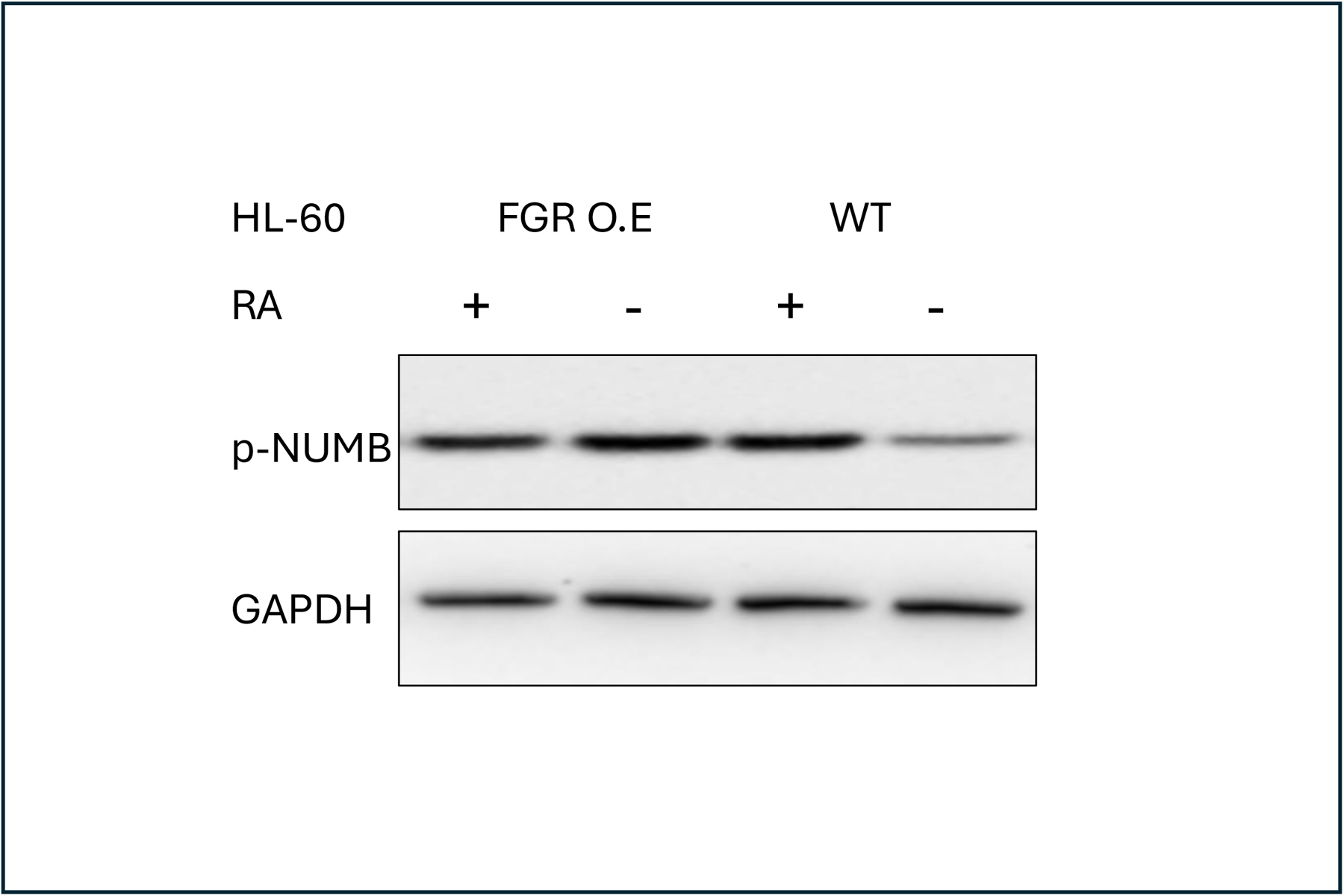
p-NUMB Western blot analysis of HL-60 wt and FGR O.E cells untreated and treated with RA. Wild-type and Fgr O.E HL-60 cells were untreated (control) or treated for 72 h with RA (1 μM) as indicated. 25 μg of lysate per lane was resolved by SDS PAGE and electro-transferred to membranes. Membrane images for each protein are cropped to show only the band of interest. For all Western blots, densitometric analysis of 3 or more biological repeats were quantified using ImageJ and shown in Figure S17.

#### FGR expression caused expression of BLR1

Expression of FGR in transfectants caused expression of BLR1 aka CXCR5. In wt cells, BLR1 is not expressed until the cells are treated with RA, which induces expression. BLR1 expression is necessary for RA-induced differentiation. Wt cells and FGR transfectants were untreated or treated with RA (1.0 microM for 72 hrs), and whole cell lysates harvested for Western blotting, probing for BLR1 (FIGURE 13). While wt cells were negative for BLR1 expression, FGR transfectants expressed BLR1. Expression of BLR in the transfectants was comparable to, if not exceeding, that in RA-treated wt cells. FGR thus induced expression of BLR1, which is known to be a necessary driver of cell differentiation in response to RA.

**Figure 13:**
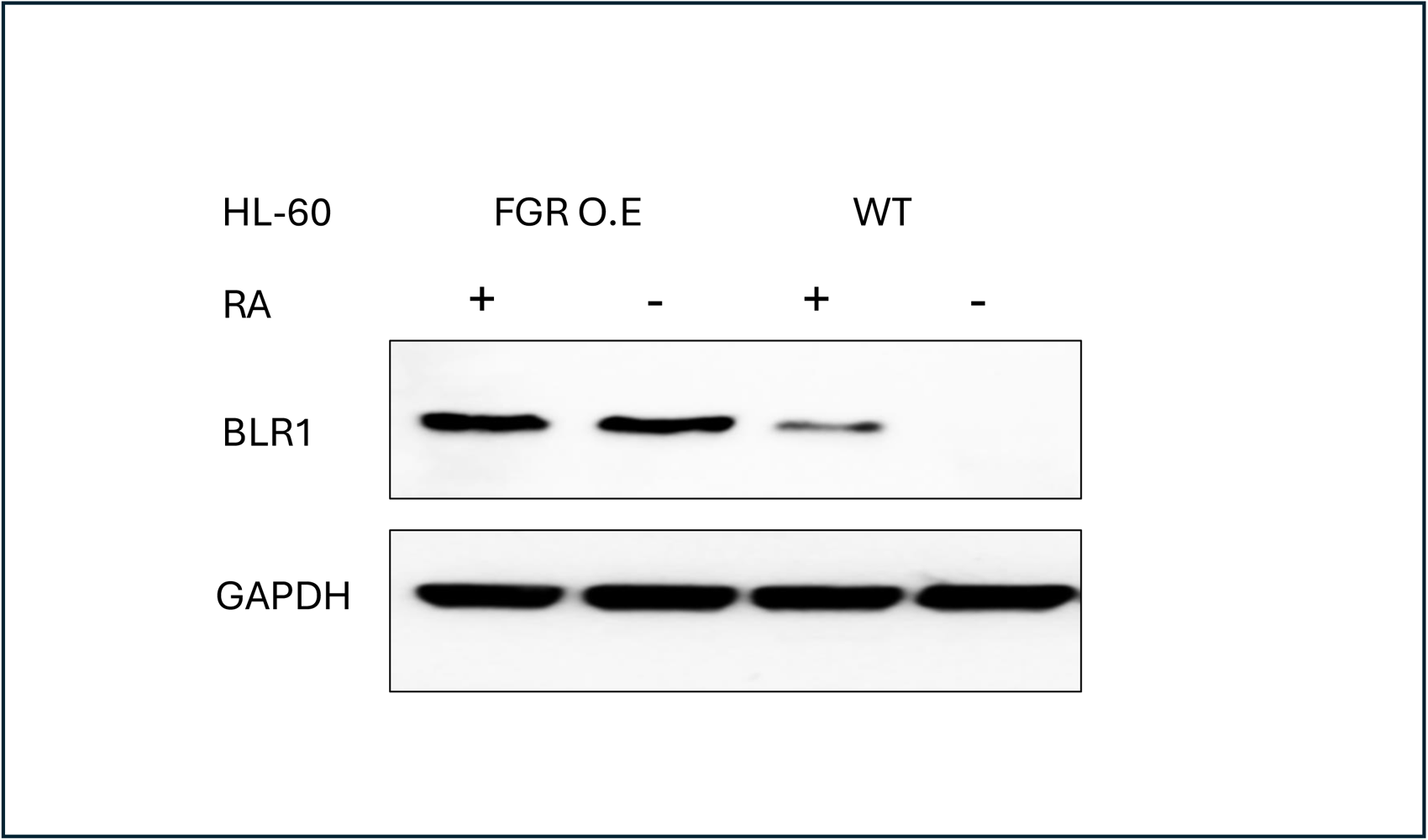
BLR1 Western blot analysis of HL-60 wt and FGR O.E cells untreated and treated with RA. Wild-type and Fgr O.E HL-60 cells were untreated (control) or treated for 72 h with RA (1 μM) as indicated. 25 μg of lysate per lane was resolved by SDS PAGE and electro-transferred to membranes. Membrane images for each protein are cropped to show only the band of interest. For all Western blots, densitometric analysis of 3 or more biological repeats were quantified using ImageJ and shown in Figure S18.

#### Ectopic expression of FGR did not rescue differentiation response of RA-resistant cells

While loss of RA-induced FGR expression is a prominent feature distinguishing RA-resistant cells created by chronic exposure to RA from their wild type parental cells, ectopic FGR expression in the resistant cells did not cause differentiation or rescue RA-induced differentiation. RA resistant variants of HL-60, called R-, were previously created by chronic exposure to RA. RA failed to induce differentiation, marked by CD38, CD11b or G1/0 enrichment. Stable transfectants expressing ectopic FGR in these cells were created as done for the wt cells (FIGURE S6). Unlike for wt parental HL-60 transfected with FGR, these transfectants failed to express CD38, CD11b or G1/0 enrichment (FIGURE S7-S9). RA-treatment also failed to elicit CD38 or CD11b expression or G1/0 enrichment. The cells were ergo still resistant despite expression of FGR, indicating that there were other dysfunctions in the resistant cells that prevented differentiation by FGR or RA, singly or in combination. There are therefore presumably collaborators with FGR that are necessary early in the process of differentiation, since even the early CD38 marker is absent. Alternatively, there is some occult regulator of signaling that blocks FGR from acting early on to start the process of differentiation. Treated with RA, the R-cells diverge from wt cells in the response of a number of signaling molecules, of which FGR was the most prominent, but any of these singly or in combination could obviously be collaborators rendered dysfunctional in the R-cells. Regardless of mechanism, FGR by itself cannot rescue resistant cells derived by chronic RA exposure. This is consistent with the notion previously put forth that resistance involves at least two steps and a number of signaling defects.

## DISCUSSION

Retinoic acid (RA) is a biologically fundamental widespread regulator of cell differentiation and proliferation in both embryonic and adult cells [2, 46]. It has been exploited as a differentiation therapy to induce differentiation and retard growth of leukemia, specifically APL, an otherwise largely incurable disease [47–50]. The present studies show that FGR causes the signaling and phenotypic shift characteristic of RA in human myeloid progenitor leukemia cells. Patient derived HL-60 human myeloid leukemia cells, which bear fidelity to a RA responsive subtype of AML [6], undergo myeloid differentiation and growth retardation in response to RA [50–52]. We now find that expression of FGR by stable transfection results in cells that are phenotypically differentiated just as if treated with RA. The transfectants express CD38, CD11b cell surface differentiation markers, the functional differentiation marker ROS that betrays the oxidative metabolism of mature myeloid cells, and the p27 CDKI responsible for G1 phase cell cycle retardation. To gain mechanistic insight, signaling known to propel RA-induced differentiation was analyzed. Previous studies have indicated that a cytosolic signalsome propels RA-induced differentiation [12–15]. Within the signalsome, RAF, FGR, LYN, SLP-76 and CBL appear as connected nodes to which other signalsome components are connected in a force-node diagram of interacting partners. RA causes increased expression of signalsome components and its activation. We now find that expression of FGR enhances expression of RAF, LYN, SLP-76 and CBL, mimicking the RA-induced up-regulation of these key signalsome components. Within the signalsome, previous studies have also indicated that VAV and NUMB appear to provide a scaffolding function. These, too, are enhanced by FGR expression. All of these signaling molecules have been previously implicated as positive regulators of RA-induced differentiation [13–19, 53], and the present studies show that FGR enhances them. A known necessary downstream consequence of the signaling in RA-treated cells is the induced expression of the BLR1 receptor, aka CXCR5, which is necessary for RA to induced differentiation [23, 24]. The present studies show that FGR expression also causes the expression of BLR1. The signaling caused by expression of FGR thus appears to be functionally competent. Hence expression of FGR causes signaling and differentiation akin to RA. FGR appears to be functioning as the single triggering event for RA-induced differentiation. It is somewhat remarkable that FGR, just a single protein by itself, can replace an agent with broad pleiotropic effects, namely RA, i.e., amongst the many effects of RA, FGR by itself was necessary and sufficient.

The present findings militate against the orthodoxy of what viral transforming genes do to enhance MAPK pathway related signaling seminal to mitogenesis. The long-standing traditional general perception of the function of FGR and SRC-family Kinases generally is to enhance MAPK pathway signaling from growth factor receptors and propel mitogenesis associated with tumorigenesis [35, 54]. The present results indicate that FGR can also do essentially the opposite; that is, propel differentiation and loss of the tumor phenotype. The effects of FGR in signal transduction, however, are varied and the present results are potentially within this broad spectrum of functions. Nevertheless, there is a paradox regarding what determines whether FGR has pro-tumor or anti-tumor effects in different contexts. One simple potential rationalization is availability of target/binding sites and stoichiometry. In the present context, accordingly there is a signalsome that binds and is activated by FGR, which presumably competes with complexes on the cytosolic domain of growth factor receptors. If the former is favored over the latter by stoichiometry or avidity, then the present phenomenon ensues. If there is no signalsome, then obviously the latter prevails. Regardless of mechanism, the present findings expand the repertoire of functions attributable to FGR.

An enigma imbedded in the present findings is how FGR causes genes controlled by RAREs to be expressed absent RA, while at the same time retaining specificity for the RA-targeted genes. For example, expression of CD38 is controlled by an RARE in the first intron [55], and expression of BLR1 is controlled by a RARE approximately 1.1 kb 5’ of the first exon [56]. Presumably, RA directed specificity must be conserved for FGR because FGR causes differentiation which depends on the proper sequential expression of specific genes. The enigma is thus how one apparently can specifically target and express RARE controlled genes without RA. While many possibilities might present themselves in the complex biology of transcriptional regulation, one possibility is that RAR and its hetero-dimerization partner, RXR, reside without any RA ligand on the promoter of a target gene and essentially license it for expression. While the mechanism is enigmatic, the present results indicate that FGR has the capability to cause expression of RARE controlled genes without RA, revealing a novel function for FGR.

Obviously, the present results indicate areas of the biology of RA and FGR where the commonly held existing signaling paradigms are incomplete. These enigmas, such as how a prototypical viral transforming agent is seminal to differentiation of leukemia cells - essentially a reversal of transformation, or how FGR causes genes controlled by RAREs to be expressed without any RA, are not as recondite if considered from the perspective of this novel paradigm for RA-induced differentiation for which the HL-60 model was the prototype. Predicated on the original finding [12] some 25 years ago that RA-induced differentiation required prolonged MAPK pathway related signaling, the now emerging paradigm is that RA causes expression of FGR, FGR binds and activates a putative signalsome in the cytosol -ostensibly at the inner membrane interface, and activation of the signalsome causes nuclear enrichment of RAF, FGR, LYN, SLP-76 and CBL, which potentially become transcription factors [15, 20]. Indeed, in support of the idea that traditionally regarded cytosolic signaling molecules can have nuclear activity, there is evidence that VAV, a GEF and adaptor, can go to the nucleus and act as a transcription factor, e.g. VAV has been found bound to NFAT in a transcriptionally active complex in one context [57]. SFKs have also been found in the nucleus and implicated in controlling chromatin condensation [58]. The prototype here is RAF (pSerine621), which binds NFATc3 downstream of the non-canonical 17bp GT box RARE of the blr1 gene, where enhanced binding due to RA is associated with transcriptional activation [22]. BLR1 expression also generates MAPK pathway signaling and is necessary for RA to induce expression of CD38, CD11b, ROS and G1/0 enrichment associated with growth retardation of differentiated cells [23, 24]. The sequence of CD38, CD11b and ROS marks the progressive phenotypic shift along the myeloid differentiation lineage. CD11b activates a CD11b/FAK/LYN/SLP-76 signaling axis to promote RA-induced HL-60 cell differentiation [13], specifically CD11b expression drives subsequent ROS and G1/0 enrichment, giving rise to the notion that the differentiation process occurs in quantized steps distinguished by successive expression of receptors. The segregation of the progression of RA-induced differentiation into two identifiable steps, namely lineage agnostic pre-commitment and then lineage specific commitment, was discovered earlier [8, 9]; and the notion of two steps recurs with the finding that resistance to RA occurred in two identifiable steps distinguishable by CD38 expression [25, 26].

In contrast to the classical transient MAPK pathway mitogenic signaling, RA-induced differentiation depends on sustained MAPK pathway activation. The ability of BLR1, CD38, and CD11b to cause MAPK pathway signaling may contribute to the prolonged MAPK signaling associated with differentiation as versus the transient signaling associated with mitogenesis. Each of these receptors can cause signaling, and their sequential expression during the process of differentiation would ostensibly result in the appearance of a long duration signal in RA-treated cells. Furthermore, each of these can activate signaling molecules that are signalsome components, eg. BLR1 can activate RAF and its consequential kinase cascade [24], CD38 can also activate RAF as well as cause phosphorylation of CBL and p85 PI3K [14], and CD11b putatively targets SLP-76 via FAK and LYN [13]; consistent with the suggestion that these signals sequentially contribute to perpetuate signalsome activation in a structurally/functionally dynamic process. A long signal originating from this process, as in the case of RA, would ergo propel differentiation whereas the short signal originating from a trans-membrane receptor, as in the case of peptide growth factors, propels mitogenesis. FGR thus starts this process by binding and activating the signalsome which results in prolonged signaling sustained by expression of induced receptors. An obvious prediction of the current data taken in the context of this paradigm is that RA must be continuously present to elicit FGR expression and drive the differentiation process, otherwise cell division will dilute the amount of FGR per cell in daughters to levels where it is no longer effective. Indeed, this anticipation was realized in previous findings showing that if RA is removed after an initial transient (24 hr) treatment, then differentiation is arrested [8, 9], however there is a brief period of memory of the previous exposure so that a subsequent RA-treatment is abbreviated for eliciting mature cells [7], but after about two division cycles this effect is lost. This paradigm provides mechanistic rationalizations for previous findings and enigmas and fundamentally expands the canonical one for how RA works.

## Abbreviations

AML: Acute myeloblastic leukemia
APC: Allophycocyanin
APL: Acute promyelocytic leukemia
c-Cbl: Casitas B-lineage Lymphoma
CD: Cluster of differentiation
CDK: Cyclin-dependent kinases
ERK: Extracellular Regulated MAP Kinase
FGR: Gardner-Rasheed feline sarcoma viral (v-fgr) oncogene homolog
GEF: Guanine nucleotide exchange factor
IRF1: Interferon Regulatory Factor 1
MAPK: Mitogen-activated protein kinase
MEK: Mitogen-activated protein kinase kinase
PML: Promyelocytic Leukemia
RAF: Rapidly Accelerated Fibrosarcoma
RAR: Retinoic acid receptor
RARE: Retinoic acid response element
RB: Retinoblastoma
RXR: Retinoid X receptor
SLP-76: SH2 domain containing leukocyte protein of 76kDa
SFK: Src family kinase

## AUTHOR CONTRIBUTIONS

NK, designed, executed and analyzed experiments, wrote manuscript; WP, created and analyzed second FGR stable transfectant; JY, provided guidance and editing; AY, designed and analyzed experiments, wrote manuscript.

## ACKNOWDGEMENT

AY is grateful for the generous support of Dr. John Babish and Paracelsian. WP is supported in part through the Cornell-City University Dual Degree program.

---

**Figure S1:**
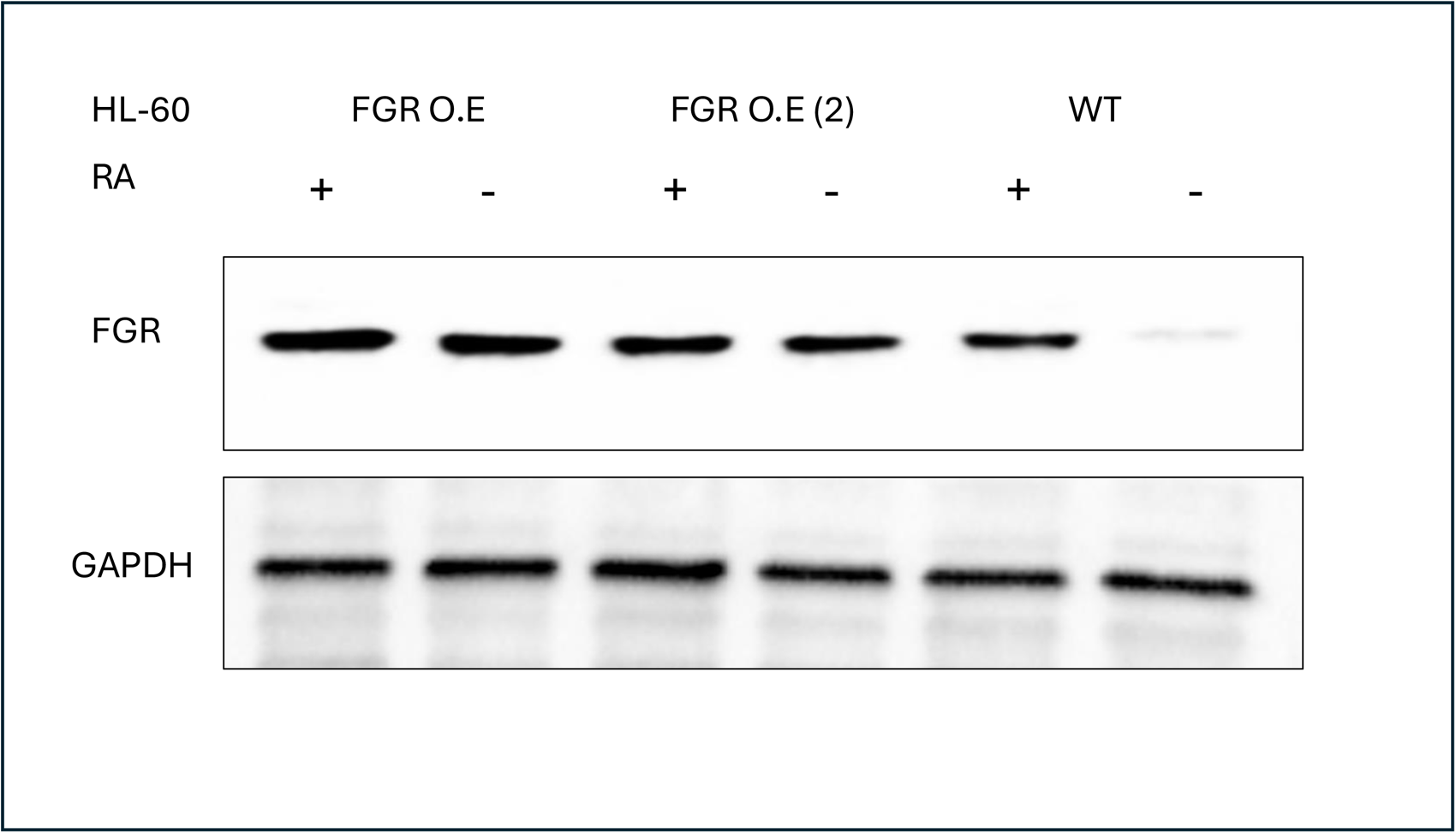
FGR Western blot analysis of HL-60 wt and FGR O.E cells untreated and treated with RA. Wild-type and Fgr O.E HL-60 cells were untreated (control) or treated for 72 h with RA (1 μM) as indicated. 25 μg of lysate per lane was resolved by SDS PAGE and electro-transferred to membranes. Membrane images for each protein are cropped to show only the band of interest. This second FGR stable transfectant was created using a lenti viral expression vector. Lenti 293T cells were seeded in 6-well plate at 70-80% confluency. On the following day, 500 uL serum free medium contains Lenti-FGR plasmid, lentivirus envelope and package plasmids pMD2.G (Addgene, #12259) and psPAX2 (Addgene, #12260) with a ratio of 2 μg : 1 μg : 2 μg was mixed with another 500 uL serum free medium contains 15 ug PEI, the mixture was allowed to stay at room temperature for 15 min; Then, the cell culture medium was replaced with the mixture above, and after incubating in the incubator for 2 h, changed with 2 mL fresh medium. After 24-30 h, the supernatant containing virus was collected, 1mL virus was added into 4 mL (density 0.1X10^6^ ml) HL-60 cells with 8 μg/mL polybrene and incubated for 6 h. Then, replacing medium with fresh 1640 medium and allowing cells growth for another 24 h. Cells were then selected in the medium containing 1ug/ml puro. This transfection method is in contrast to the original FGR stable transfectant; indicating that effects of FGR transfection were ergo not vector specific/dependent.

**Figure S2:**
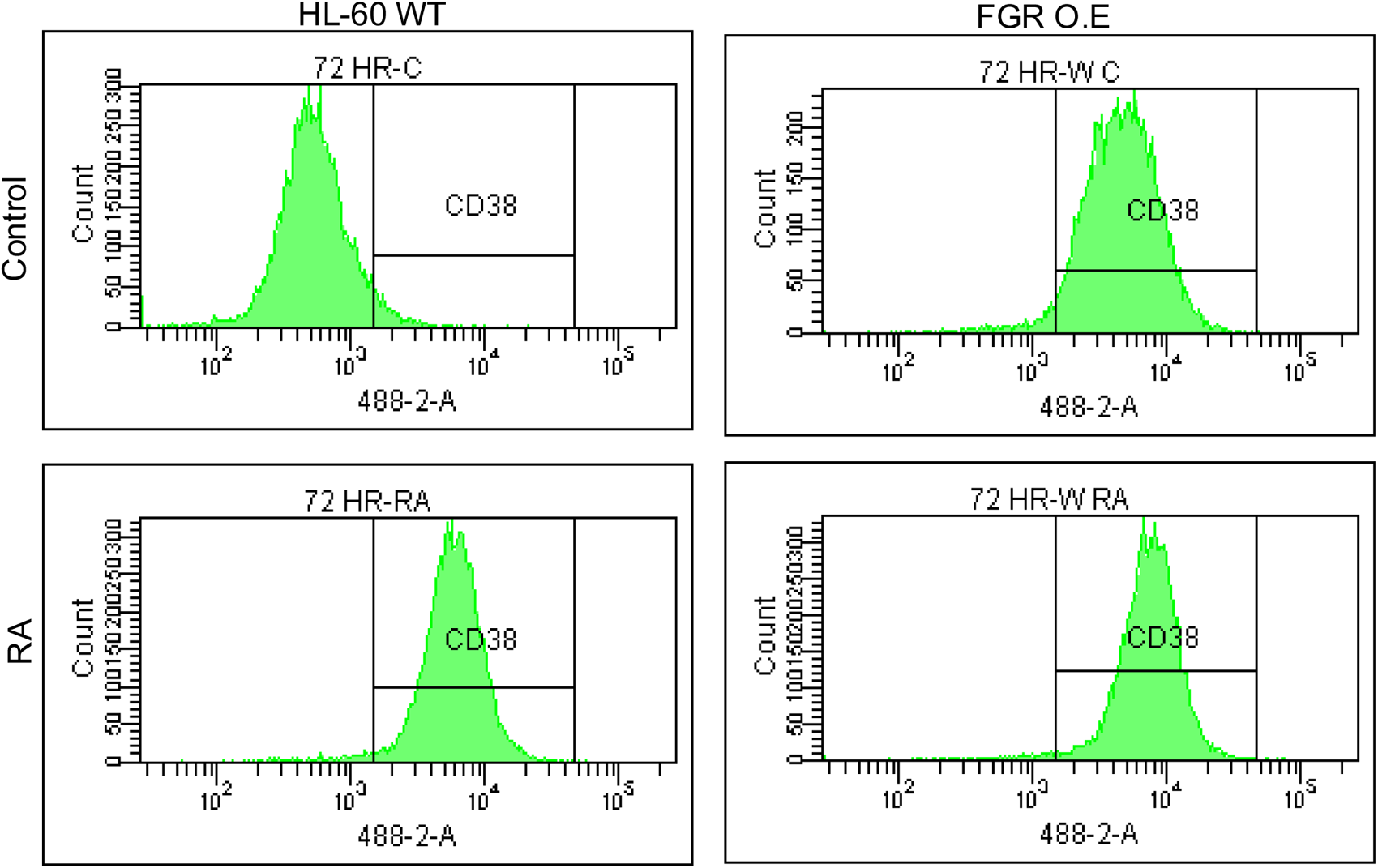
Phenotypic cell surface differentiation marker analysis of HL-60 wt and FGR O.E cells untreated and treated with RA. HL-60 cells were cultured in the absence (control) or presence of 1 μM RA as indicated. CD38 expression was assessed by flow cytometry following 72 h treatment period. Gating to discriminate positive cells was set to exclude 95% of untreated controls.

**Figure S3:**
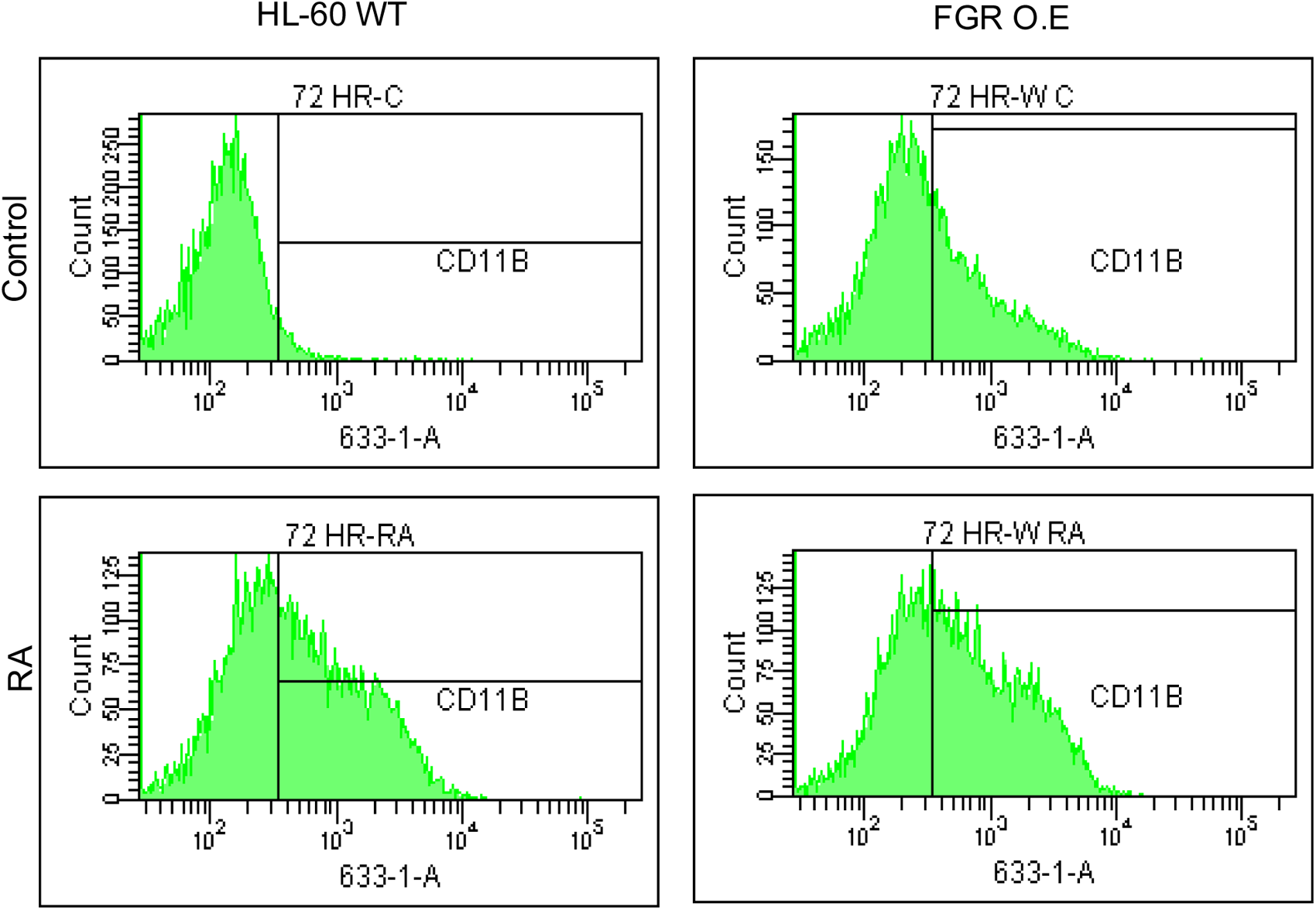
Phenotypic cell surface differentiation marker analysis of HL-60 wt and FGR O.E cells untreated and treated with RA. CD11b expression was assessed by flow cytometry after 72 h treatment periods Gating to discriminate positive cells was set to exclude 95% of untreated controls.

**Figure S4:**
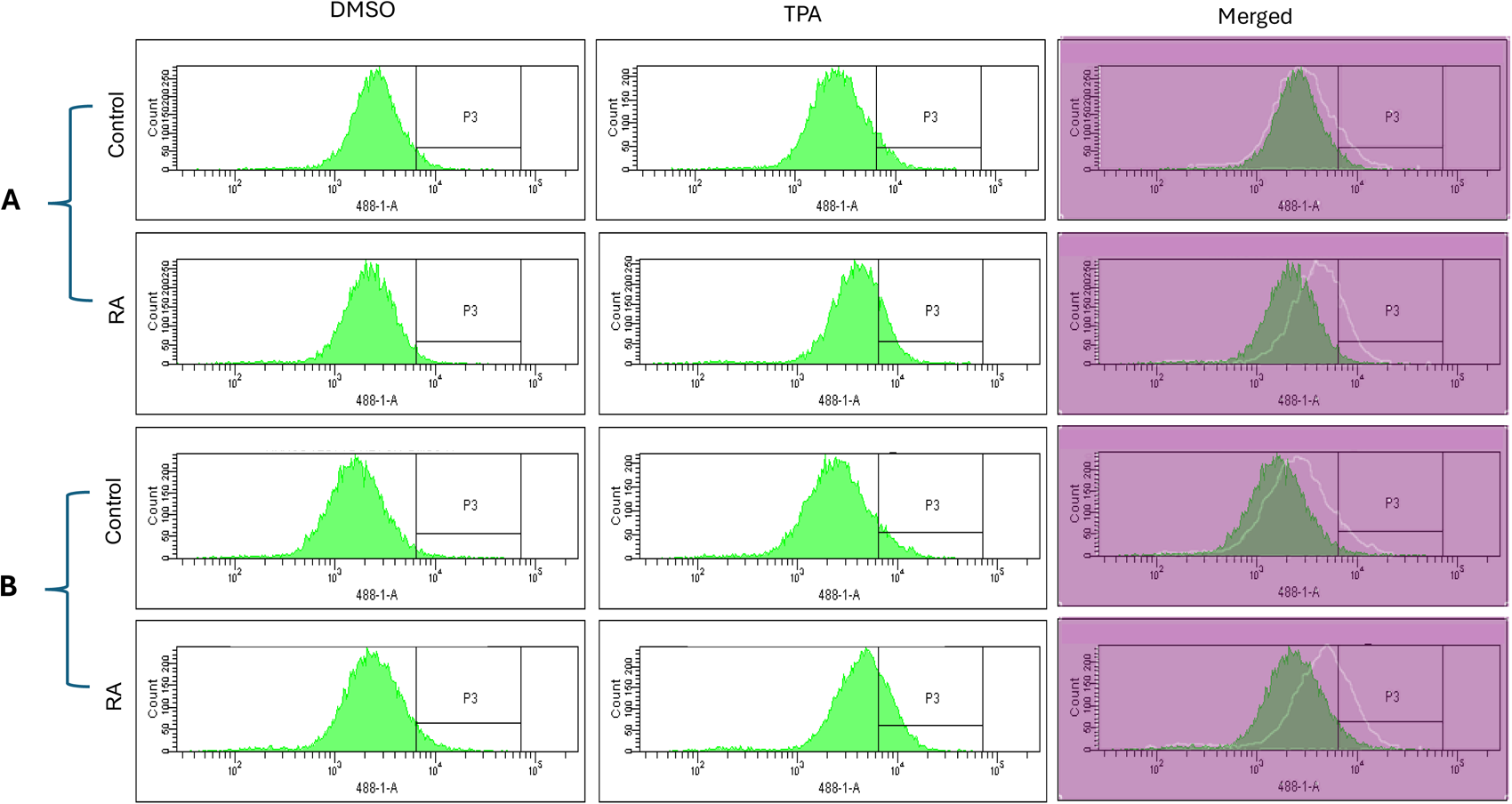
Functional differentiation marker analysis of HL-60 wt and FGR O.E cells untreated and treated with RA measured by TPA-induced respiratory burst. (**A**) HL-60 WT (parental wildtype) cells were cultured in the absence (control) or presence of 1 μM RA as indicated. (**B**) FGR O.E cells were cultured in the absence or presence of 1 μM RA as indicated. Respiratory burst was analyzed by measuring reactive oxygen species (ROS) production by flow cytometry using the 2′,7′-dichlorofluorescein (DCF) assay for DMSO carrier control and TPA induced cells. Gates shown in the histograms were set to exclude 95% of the DMSO-treated control population (carrier control) for each culture condition. For each of the 4 cases, WT and FGR that were control and RA-treated, TPA-treated samples show induced ROS. Inducible ROS production is betrayed by the shift in the TPA histogram compared to the DMSO histogram shown in the merged histogram.

**Figure S5:**
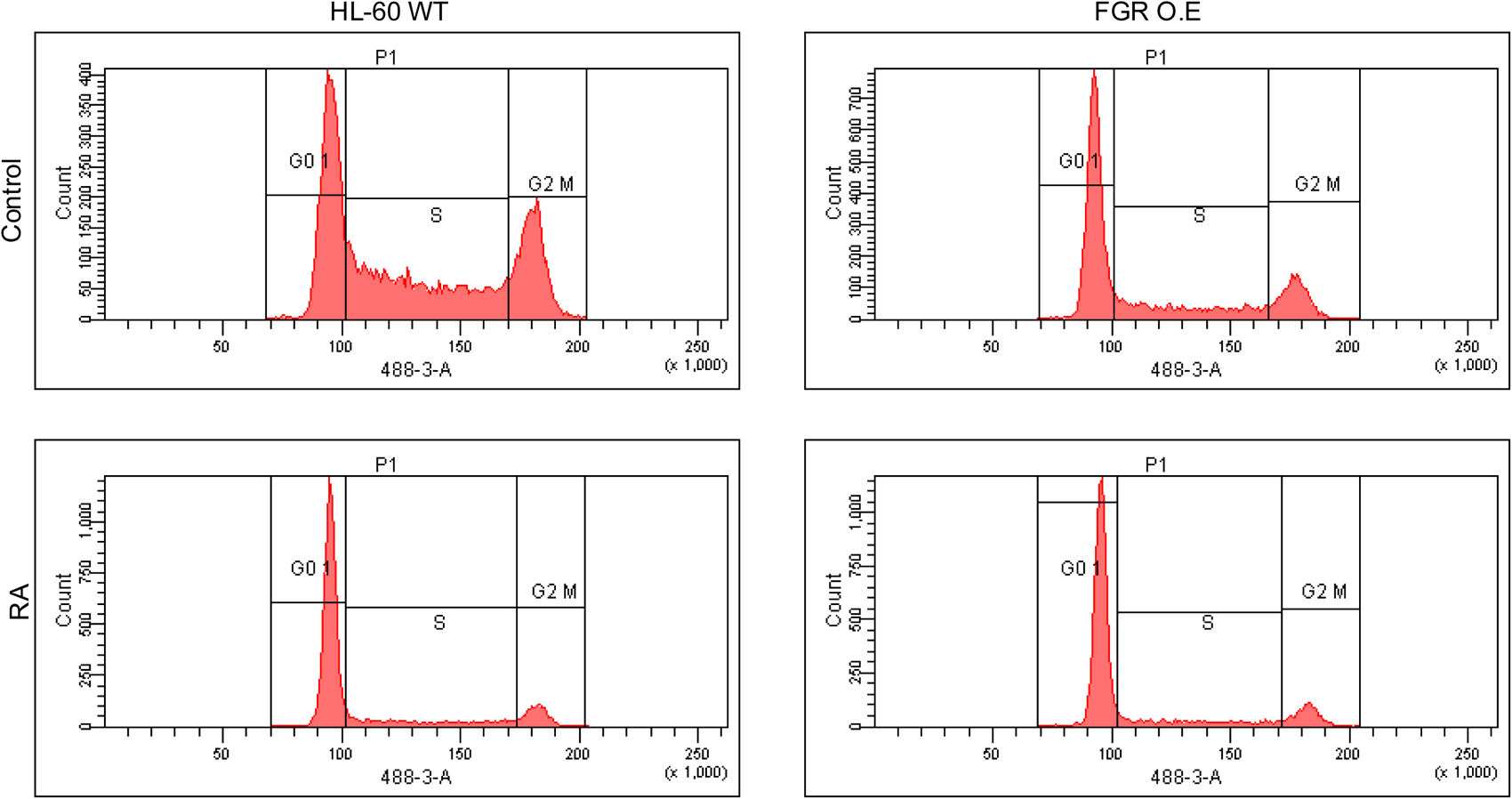
Cell cycle analysis of HL-60 WT and FGR O.E cells. DNA histograms show FGR transfectants were enriched for relative number of G1/0 cells compared to the wt cells. Wild-type and FGR O.E cell lines were cultured for 72 h without (untreated control) or with 1 μM RA as indicated. Cell cycle distribution showing the percentage of cells in G1/G0 was analyzed using flow cytometry with propidium iodide staining at 72 h. Gates define the G1, S, and G2/M subpopulations (left to right). G1/0 arrest is indicated by an increase in the G1 peak for FGR O.E compared to wt cells. The G1/0 enrichment of FGR stable transfectants compared to WT cells is associated with slower population growth of these FGR stable transfectants, as well as the original transfectants, as expected. Histogram shows percentage of cells in each phase.

**Figure S6:**
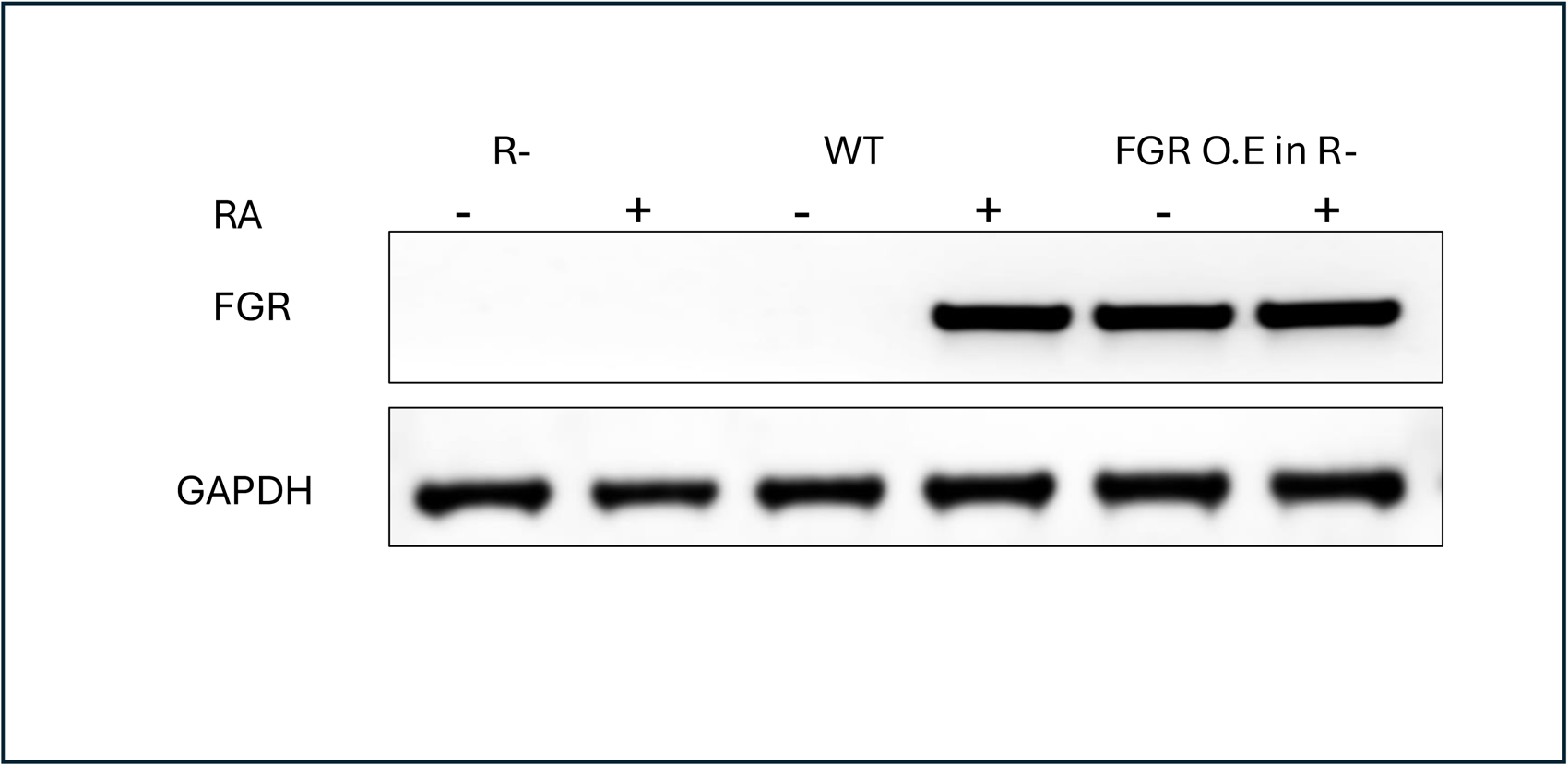
FGR Western blot analysis of HL-60 WT, HL-60 R-resistant cells and FGR O.E in R-cells untreated and treated with RA. Wild-type and FGR O.E HL-60 cells were untreated (control) or treated for 72 h with RA (1 μM) as indicated. 25 μg of lysate per lane was resolved by SDS PAGE and electro-transferred to membranes. Membrane images for each protein are cropped to show only the band of interest.

**Figure S7:**
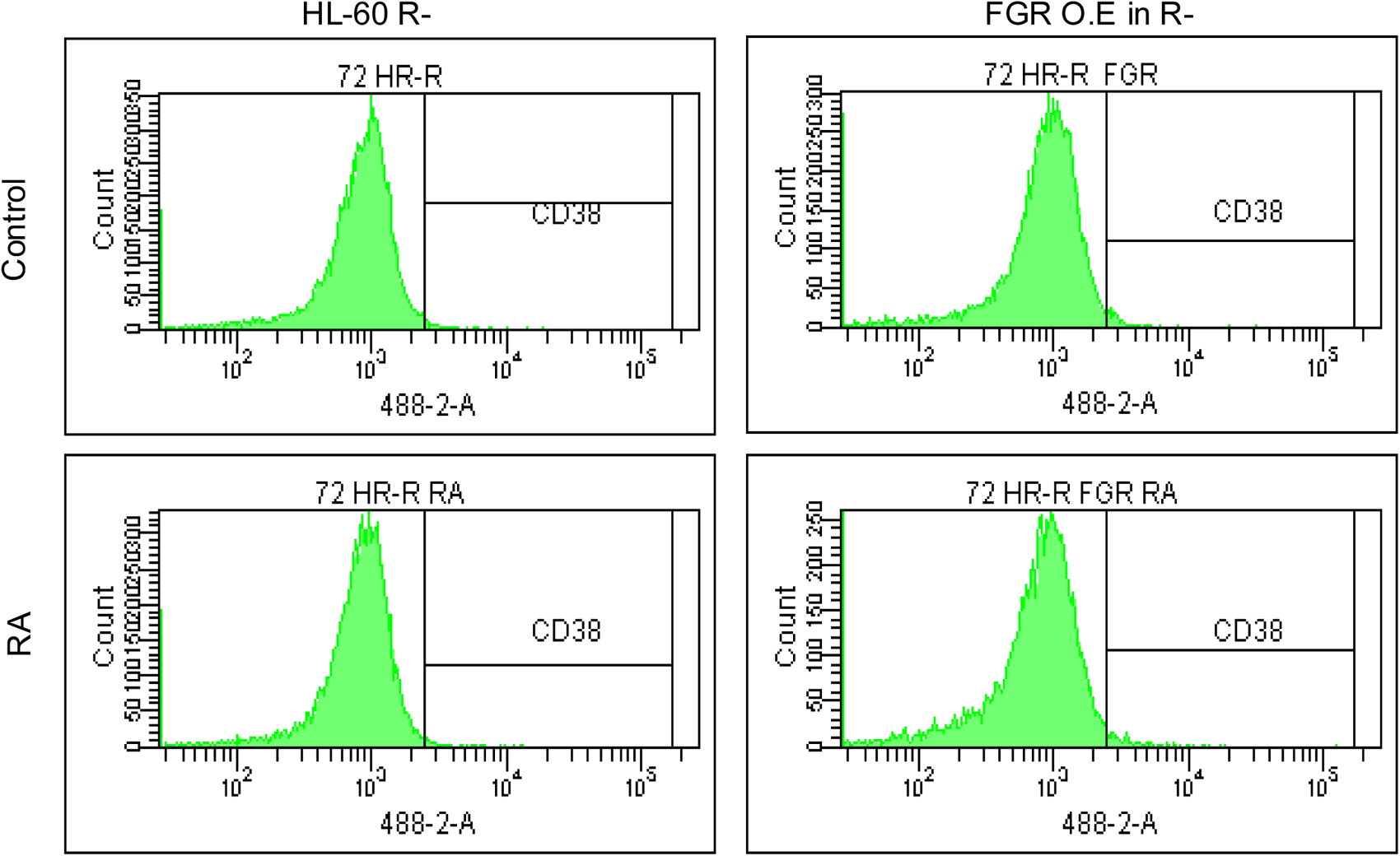
Phenotypic cell surface differentiation marker analysis of HL-60 R-resistant cells and FGR O.E in R-cells untreated and treated with RA. HL-60 cells were cultured in the absence (control) or presence of 1 μM RA as indicated. CD38 expression was assessed by flow cytometry following 72 h treatment period. Gating to discriminate positive cells was set to exclude 95% of untreated controls.

**Figure S8:**
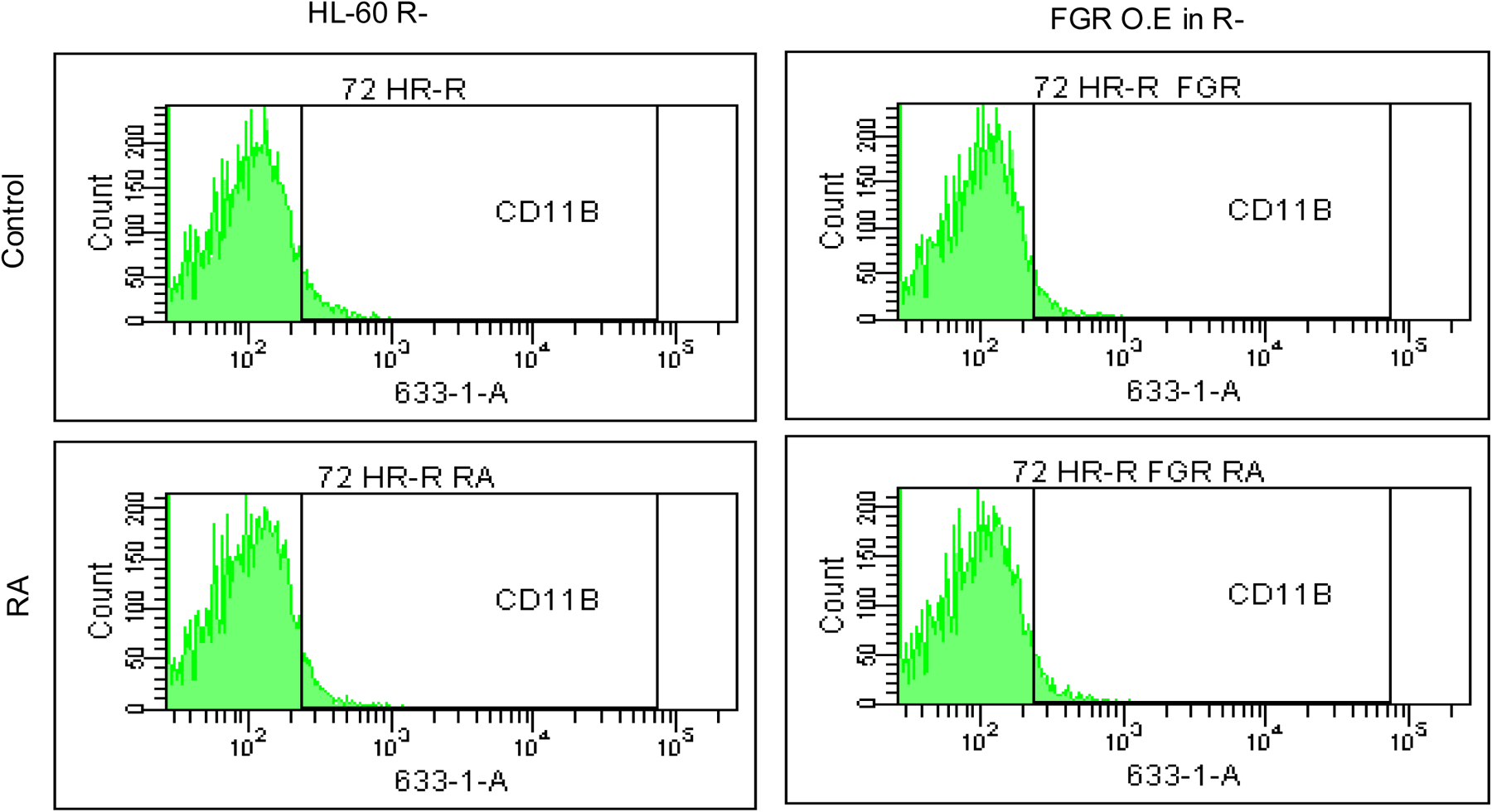
Phenotypic cell surface differentiation marker analysis of HL-60 R-resistant cells and FGR O.E in R-cells untreated and treated with RA. CD11b expression was assessed by flow cytometry after 72 h treatment periods Gating to discriminate positive cells was set to exclude 95% of untreated controls.

**Figure S9:**
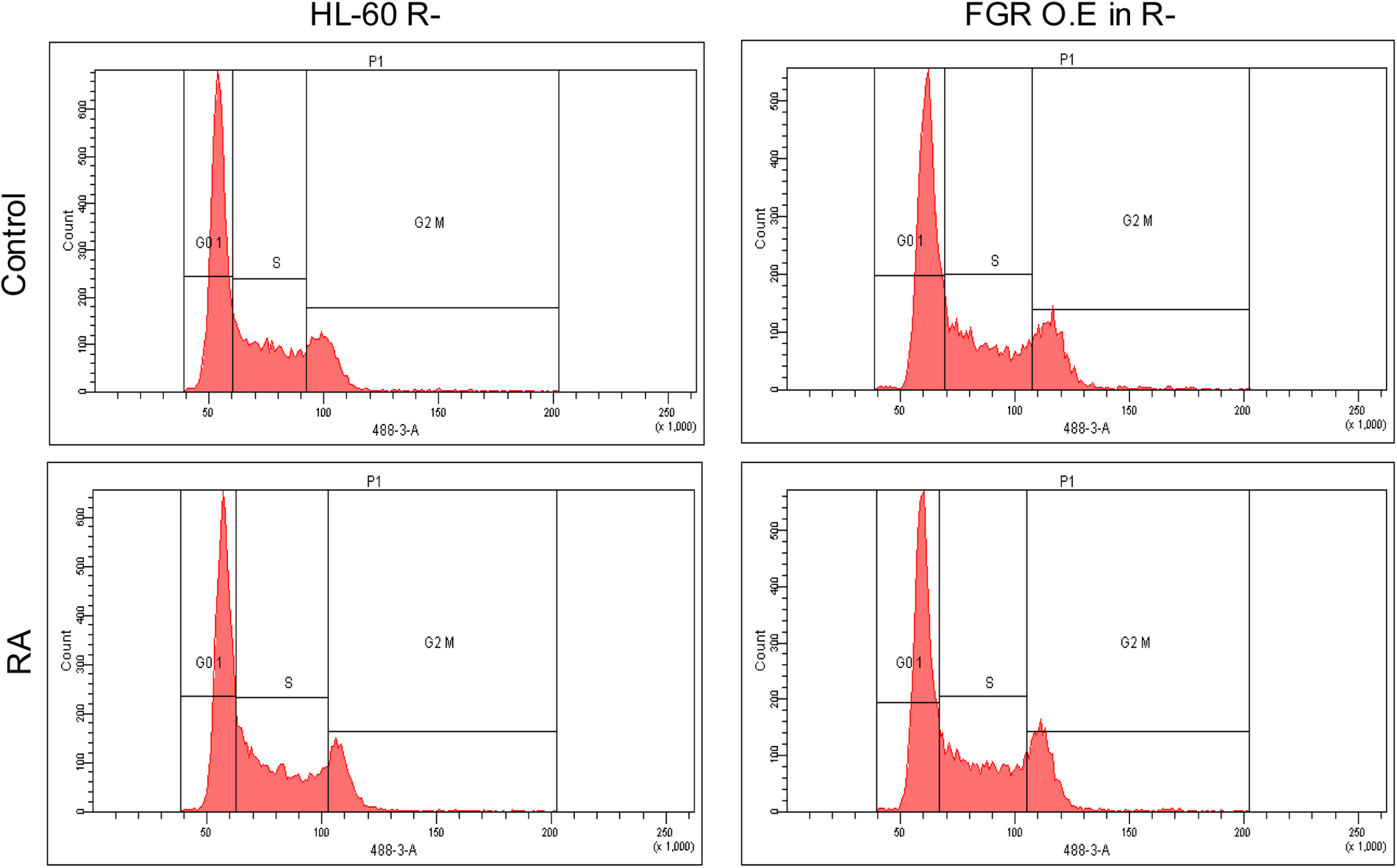
Cell cycle analysis of HL-60 R-resistant cells and FGR O.E in R-cells. HL-60 R- and FGR O.E in R-cell lines were cultured for 72 h without (untreated control) or with 1 μM RA as indicated. Cell cycle distribution showing the percentage of cells in G1/G0 was analyzed using flow cytometry with propidium iodide staining at 72 h. Gates define the G1, S, and G2/M subpopulations (left to right). Lack of G1/0 arrest is indicated by a lack of increase in the G1 peak for HL-60 R- and FGR O.E in R-control and RA treated cells.

**Figure S10:**
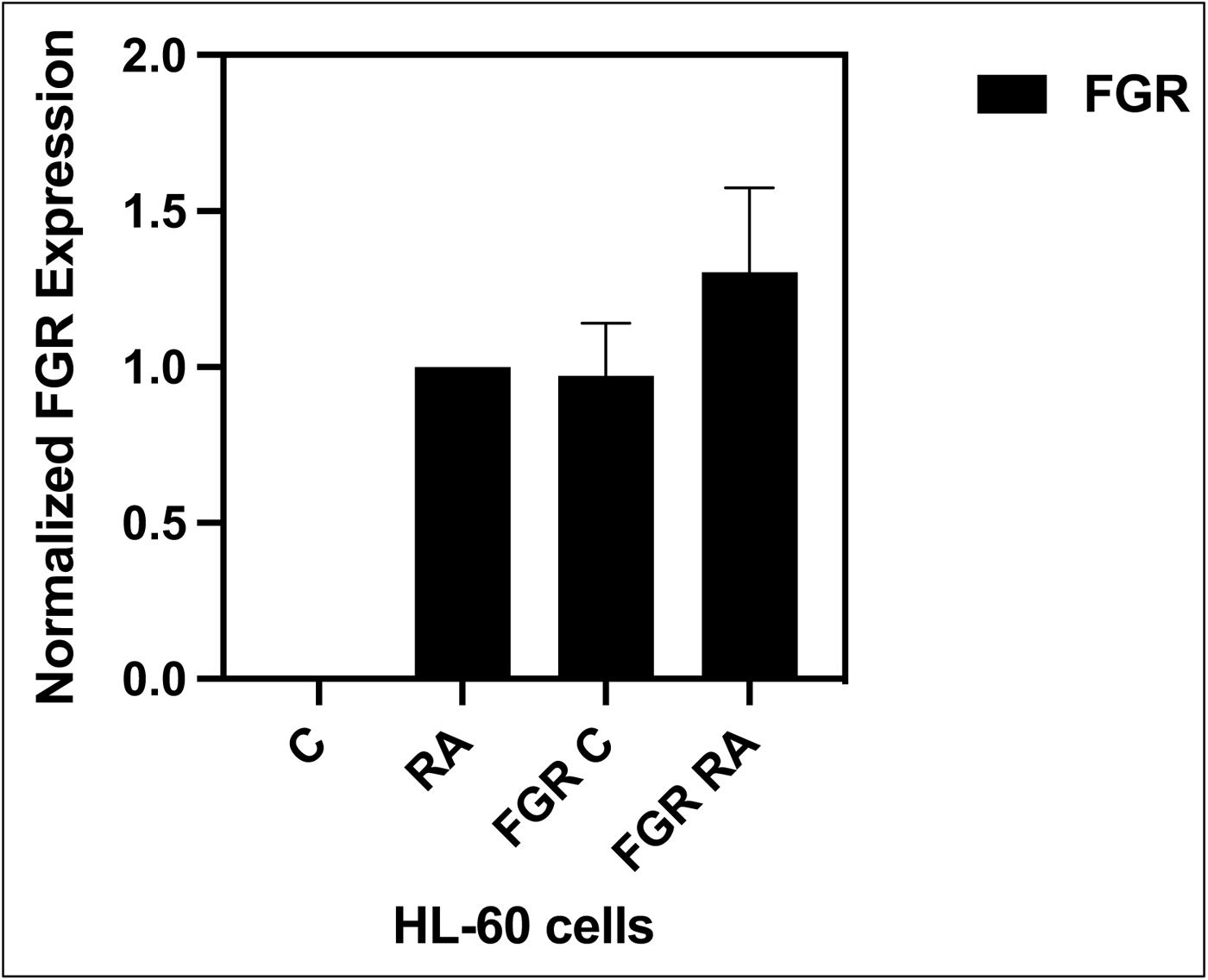
The FGR histogram. shows normalized densitometric values of the FGR western blot analysis of HL-60 Wt and FGR O.E cells untreated and treated with RA. (*n* = 3). Error bars indicate SEM.

**Figure S11:**
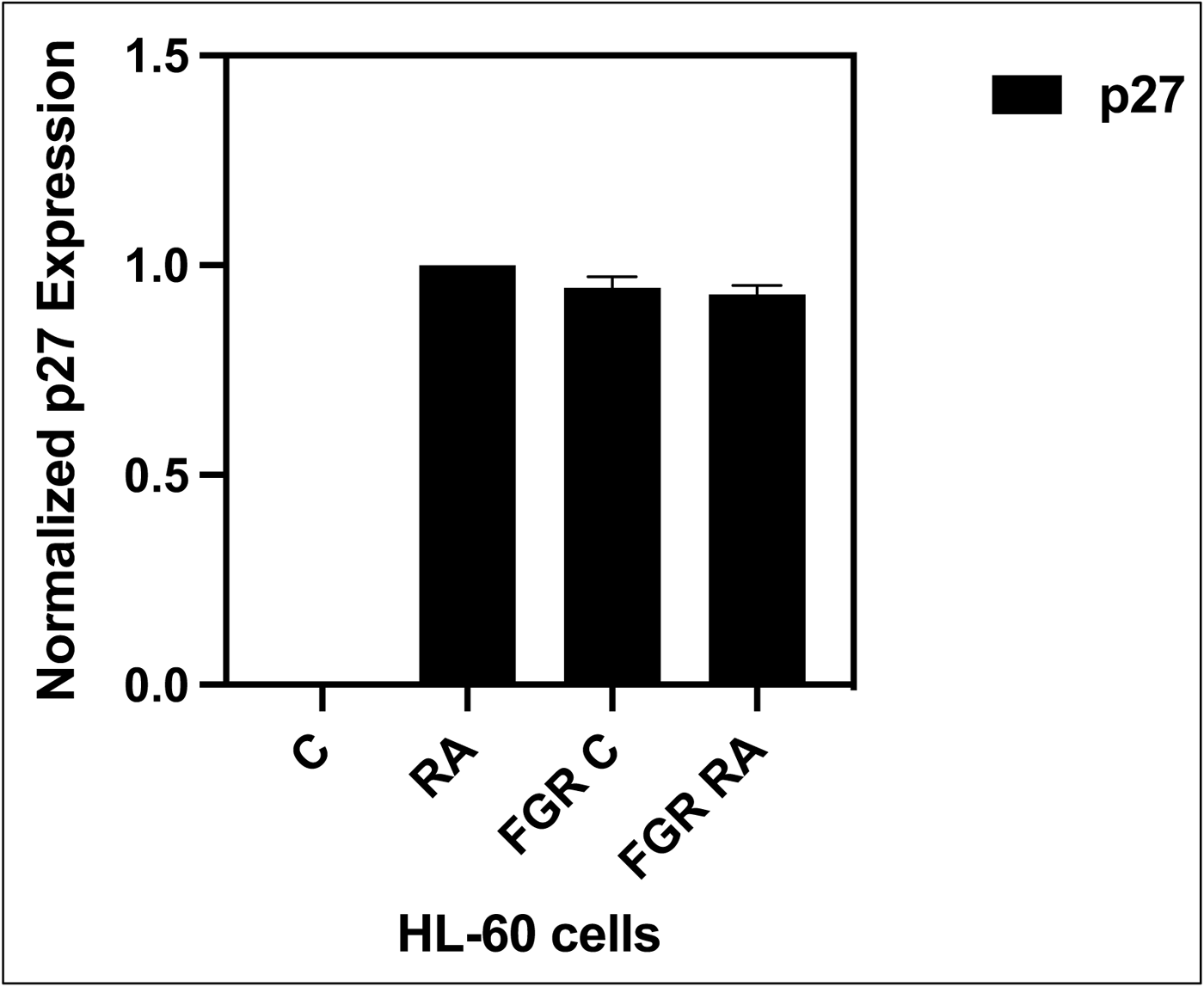
The p27 histogram. shows normalized densitometric values of the p27 western blot analysis of HL-60 Wt and FGR O.E cells untreated and treated with RA. (*n* = 3). Error bars indicate SEM.

**Figure S12:**
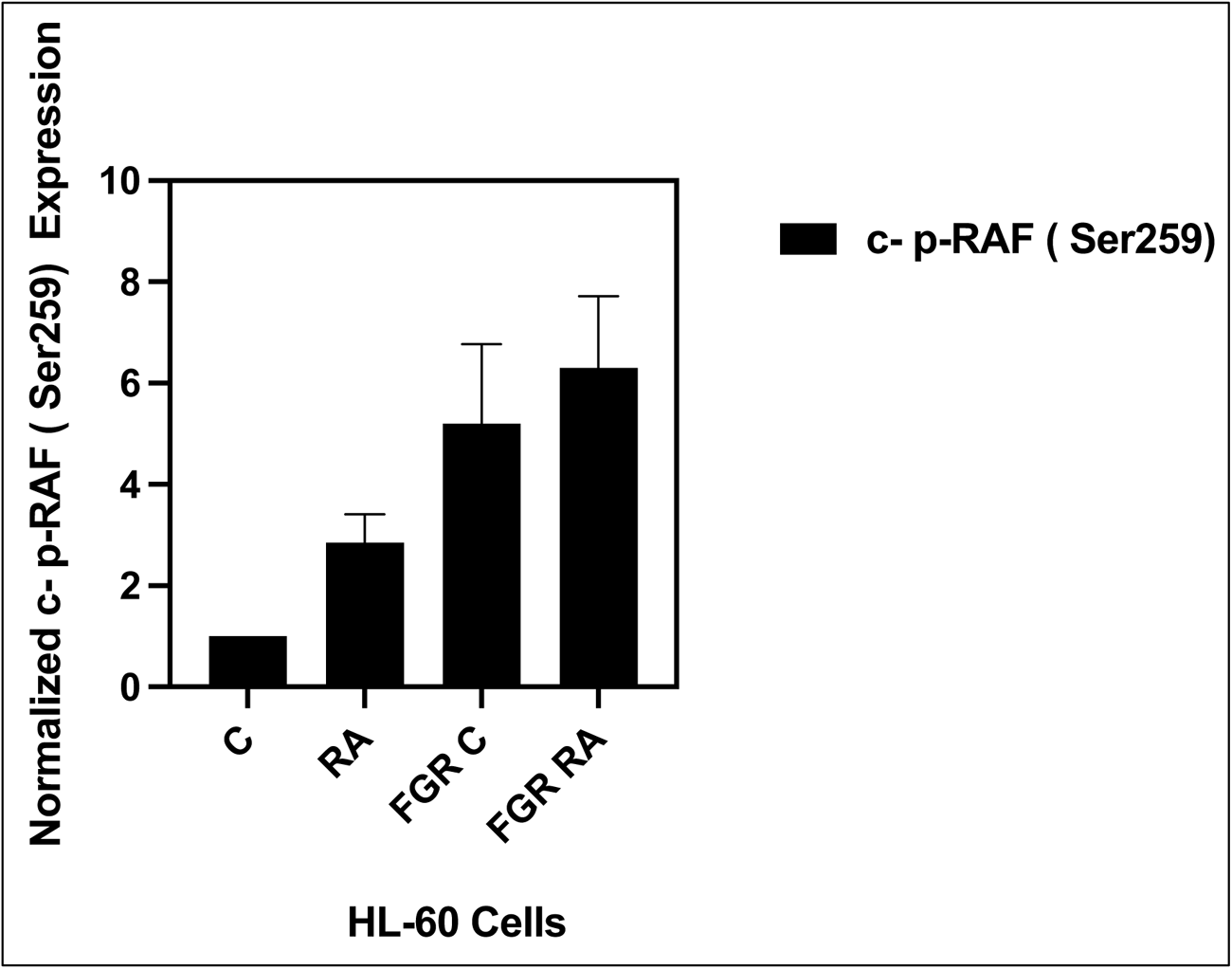
p-c-RAF (Ser259) histogram. shows normalized densitometric values of the p-c-RAF (Ser259) western blot analysis of HL-60 wt and FGR O.E cells untreated and treated with RA. (*n* = 3). Error bars indicate SEM.

**Figure S13:**
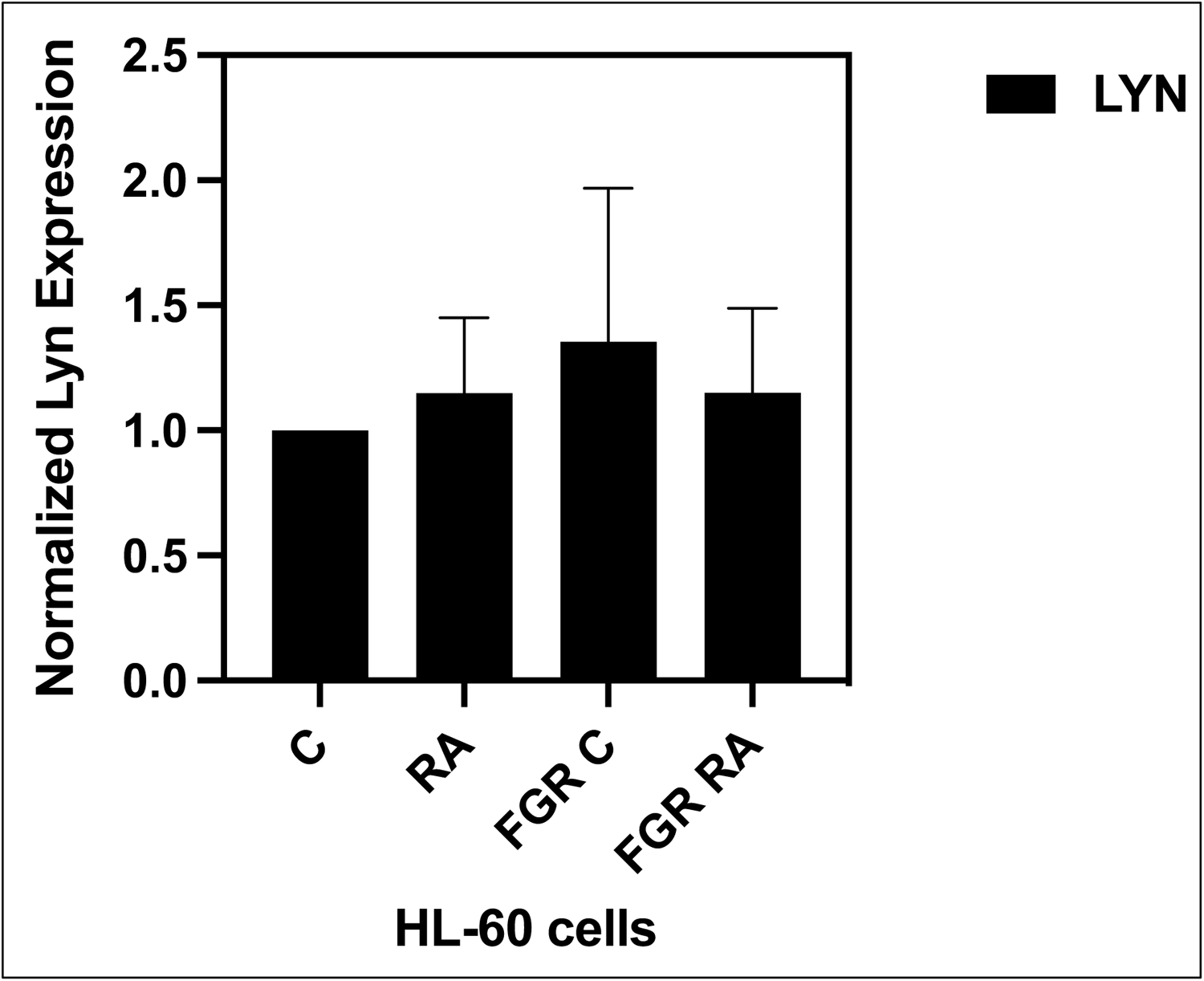
The LYN histogram. shows normalized densitometric values of the LYN western blot analysis of HL-60 wt and FGR O.E cells untreated and treated with RA. (*n* = 3). Error bars indicate SEM.

**Figure S14:**
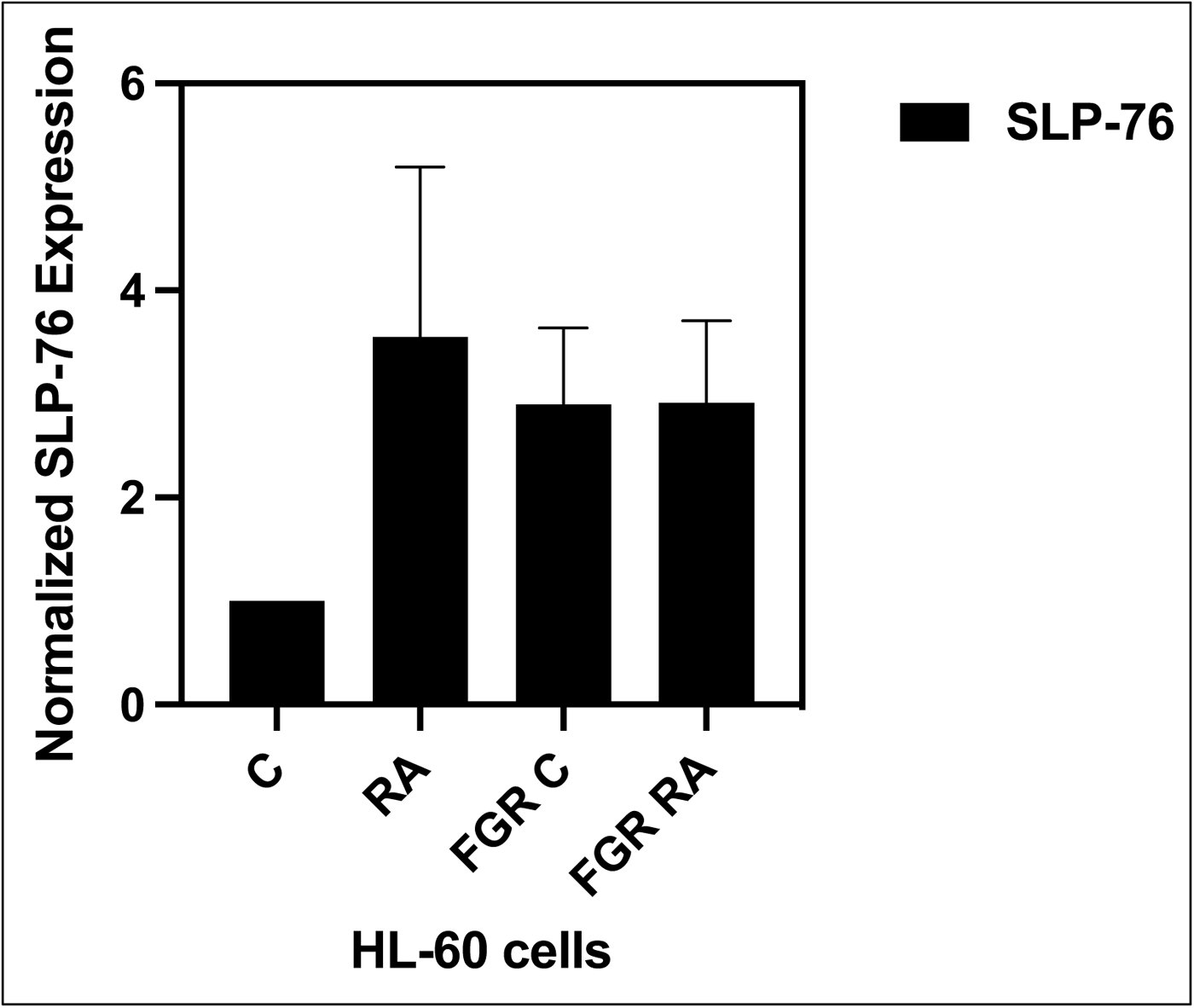
The SLP-76 histogram. shows normalized densitometric values of the SLP-76 western blot analysis of HL-60 wt and FGR O.E cells untreated and treated with RA. (*n* = 3). Error bars indicate SEM.

**Figure S15:**
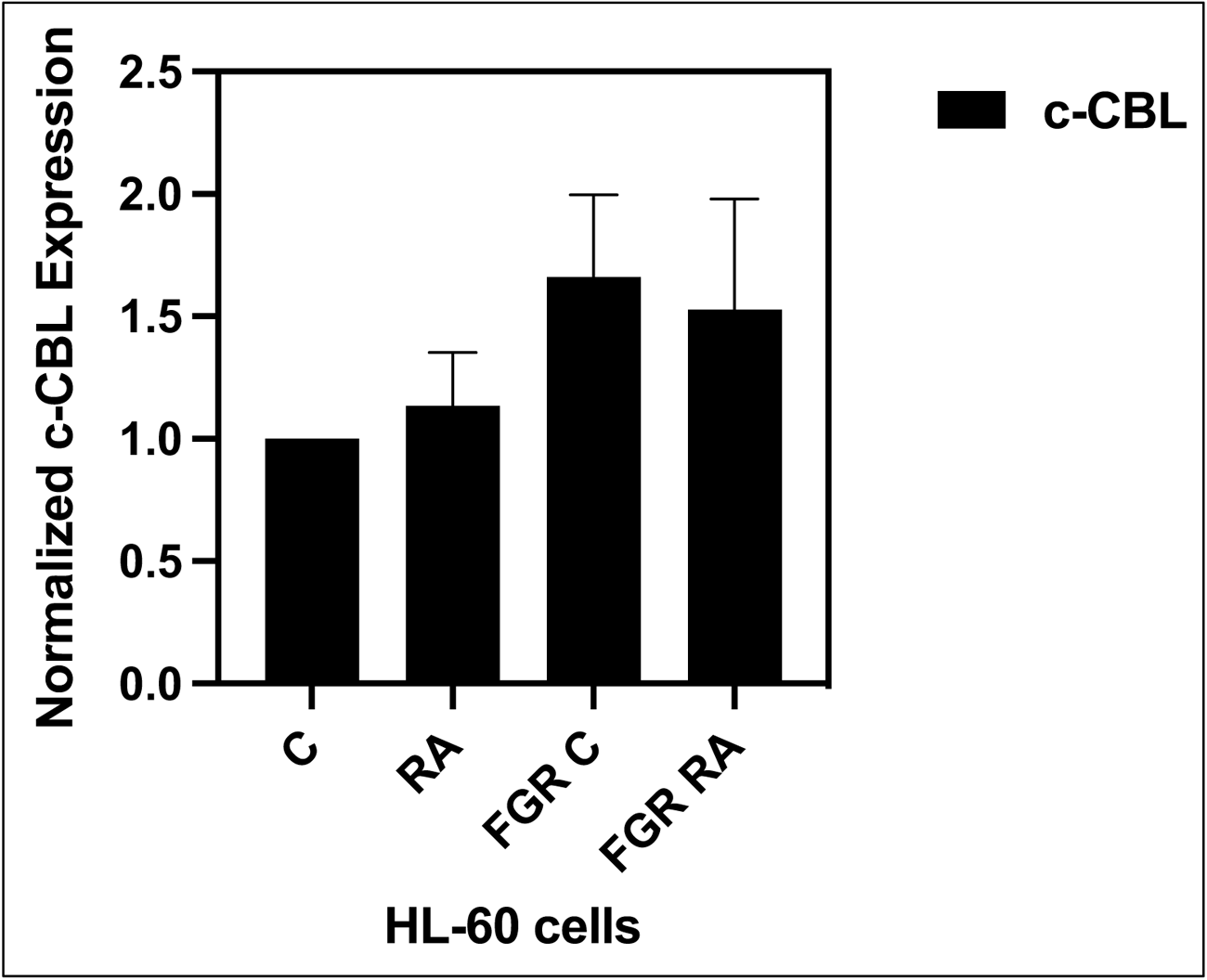
The c-CBL histogram. shows normalized densitometric values of the c-CBL western blot analysis of HL-60 wt and FGR O.E cells untreated and treated with RA. (*n* = 3). Error bars indicate SEM.

**Figure S16:**
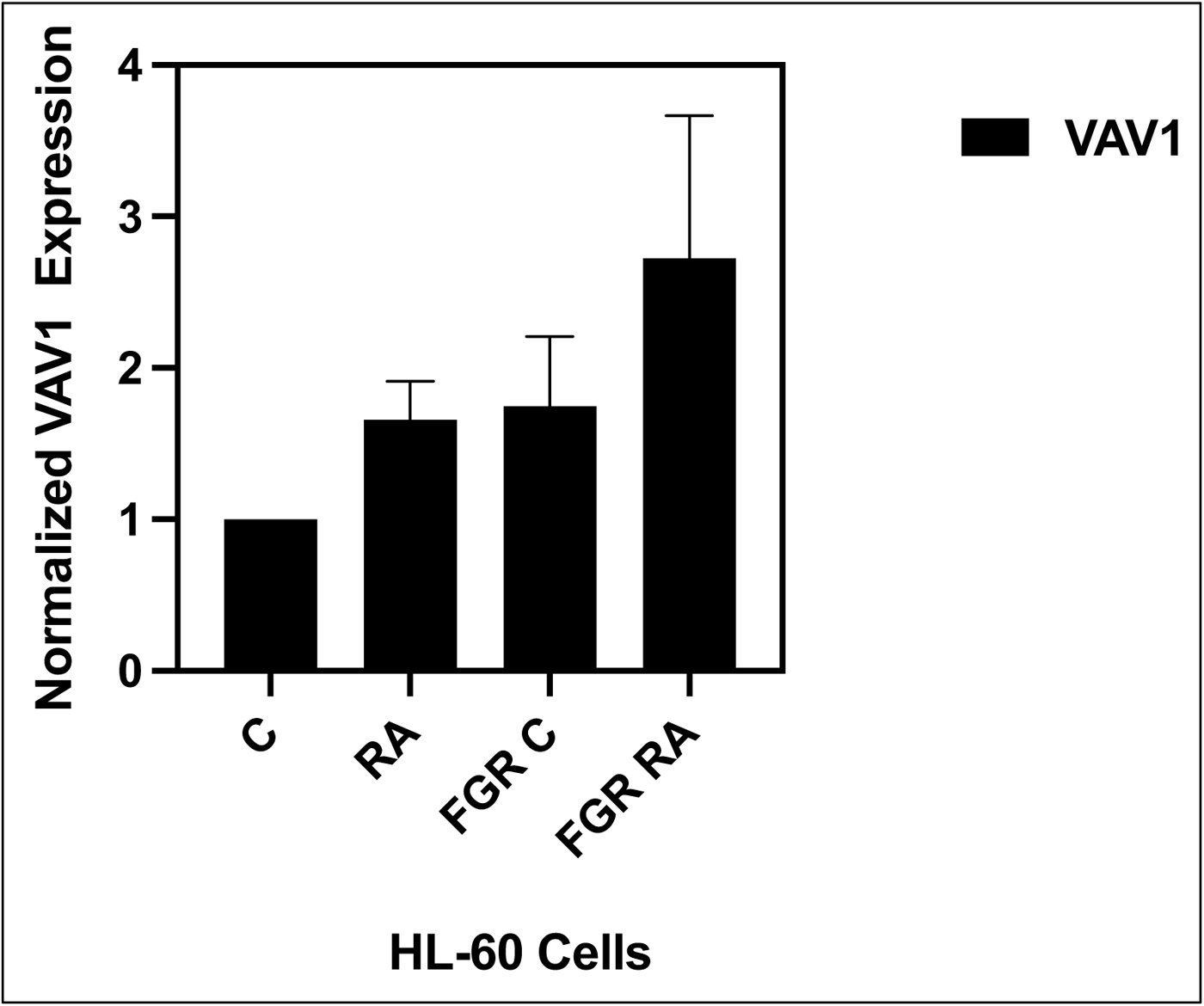
The VAV1 histogram. shows normalized densitometric values of the VAV1 western blot analysis of HL-60 wt and FGR O.E cells untreated and treated with RA. (*n* = 3). Error bars indicate SEM.

**Figure S17:**
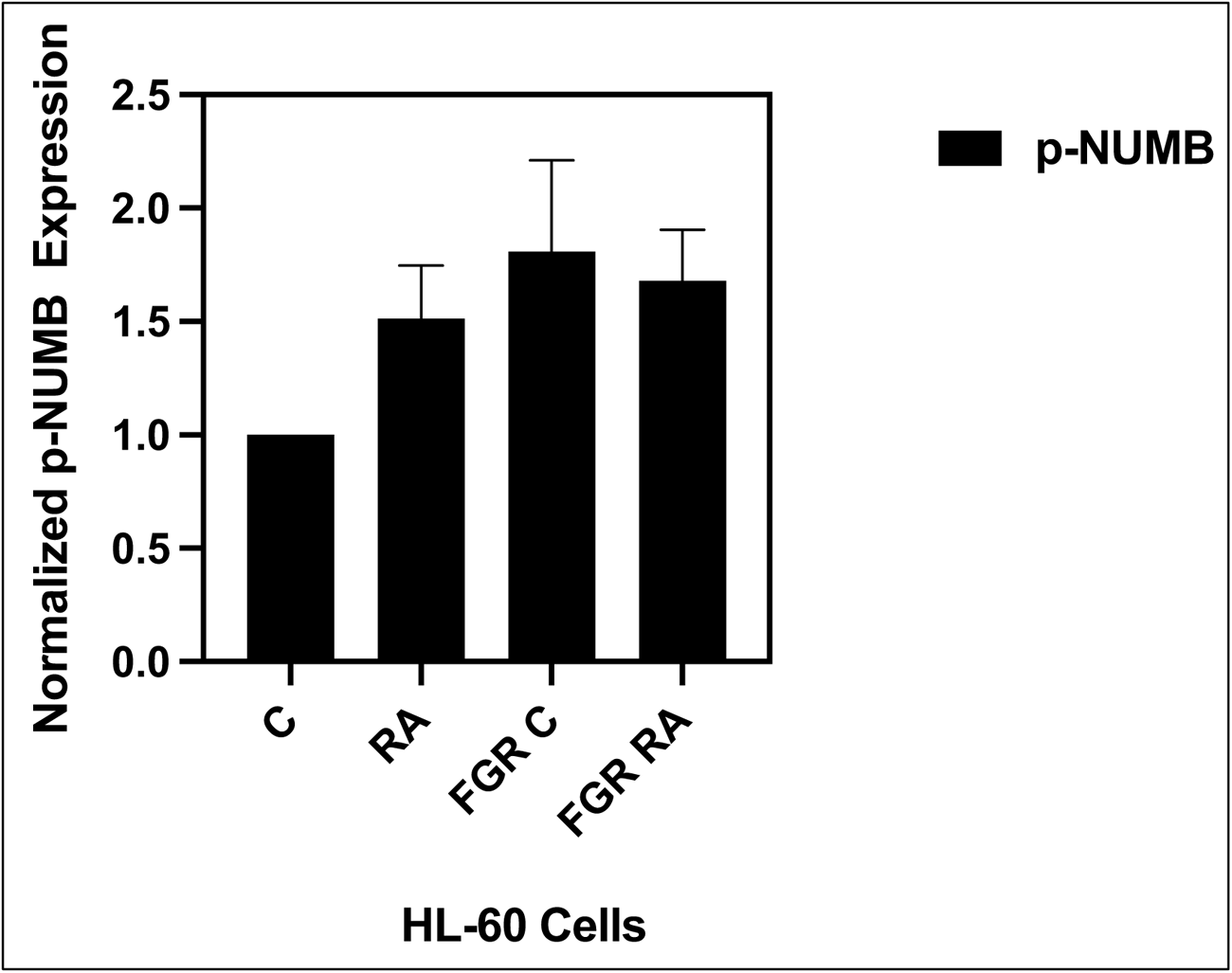
The p-Numb histogram. shows normalized densitometric values of the p-Numb western blot analysis of HL-60 wt and FGR O.E cells untreated and treated with RA. (*n* = 3). Error bars indicate SEM.

**Figure S18:**
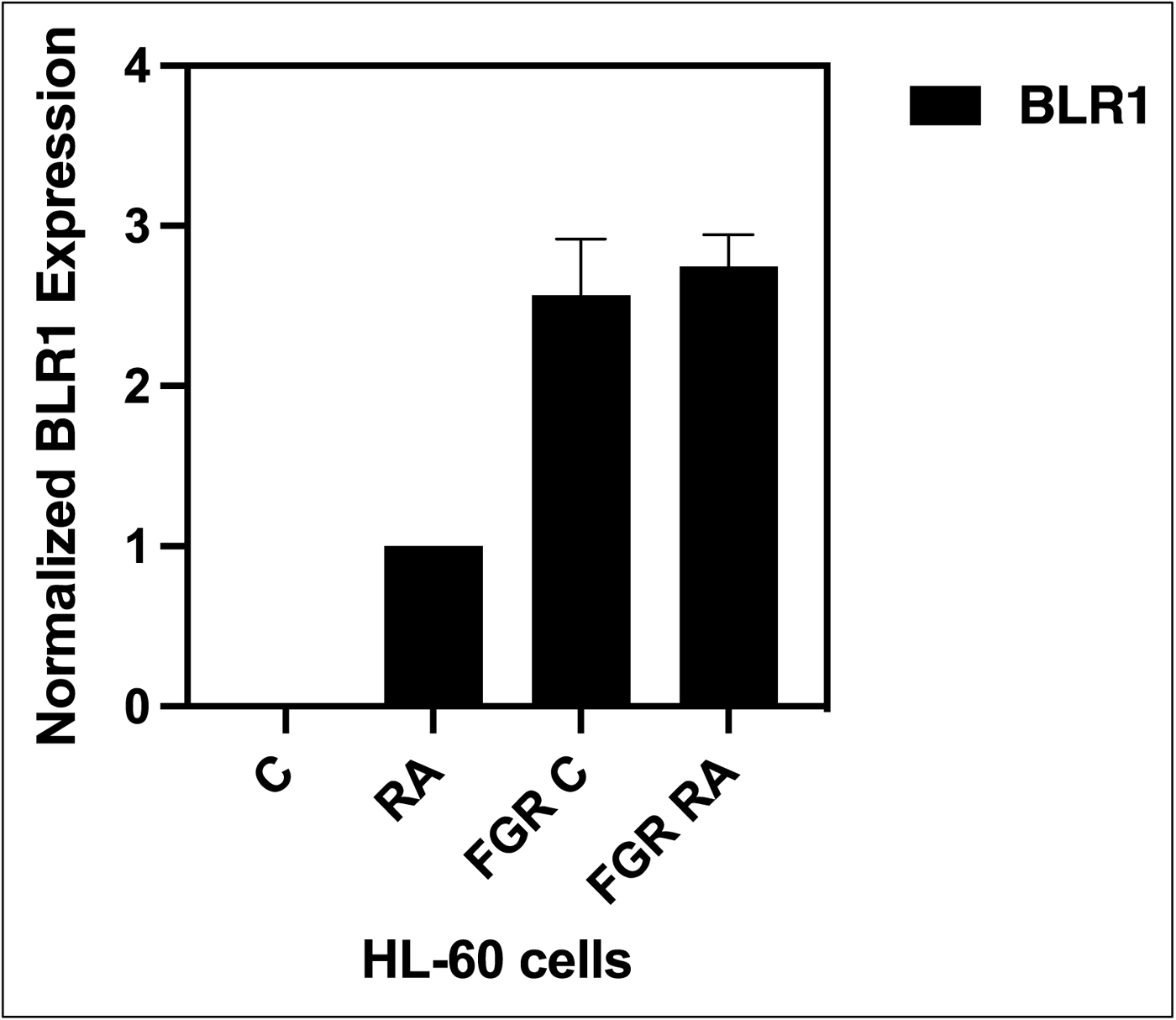
The BLR1 histogram. shows normalized densitometric values of the BLR1 western blot analysis of HL-60 wt and FGR O.E cells untreated and treated with RA. (*n* = 3). Error bars indicate SEM.

**Figure S19:**
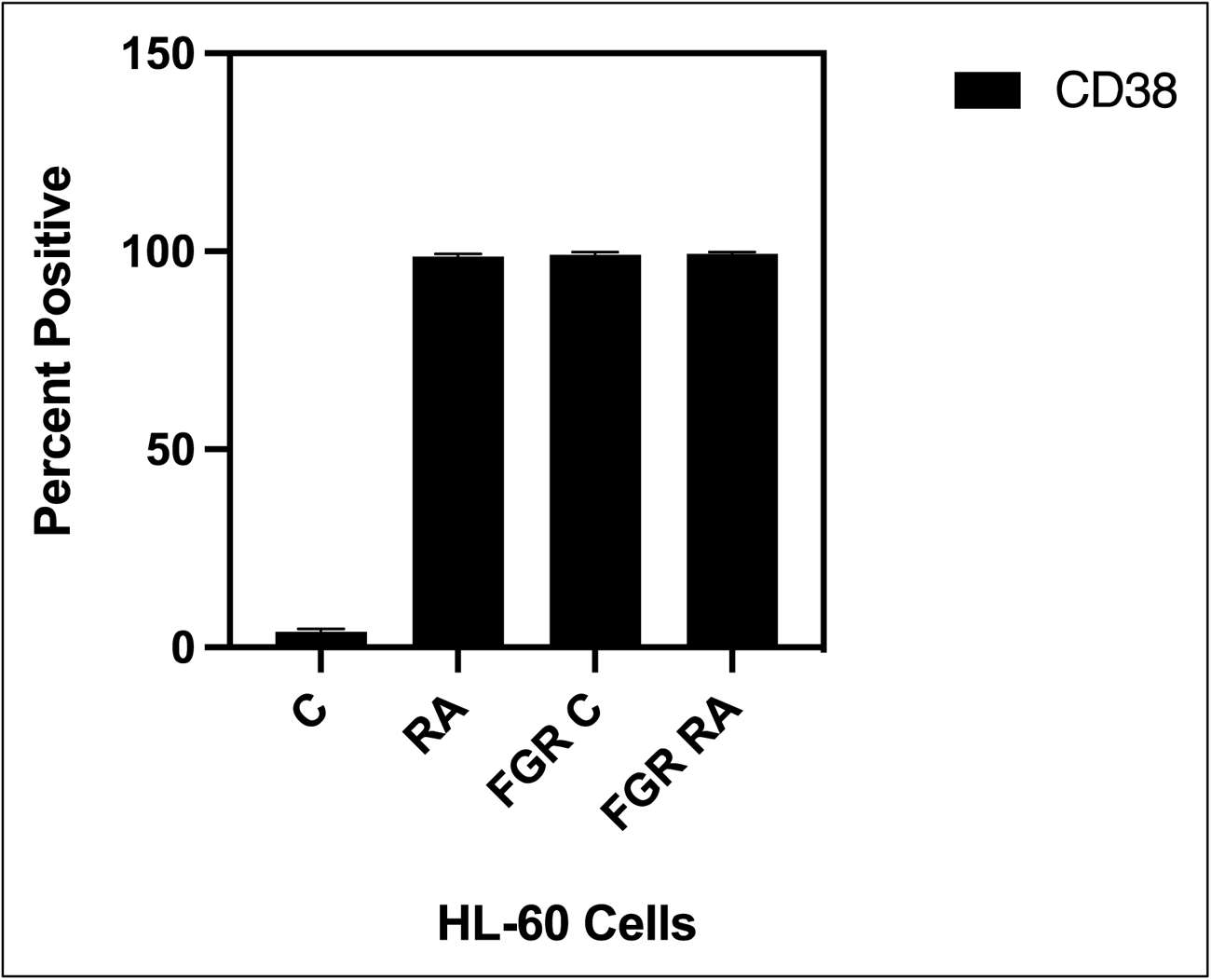
The CD38 expression histogram. Percentage of cells positive for CD38 expression at 72 h (*n* = 3). Error bars indicate SEM comparing RA-treated HL-60 wt samples to RA FGR O.E cells samples.

**Figure S20:**
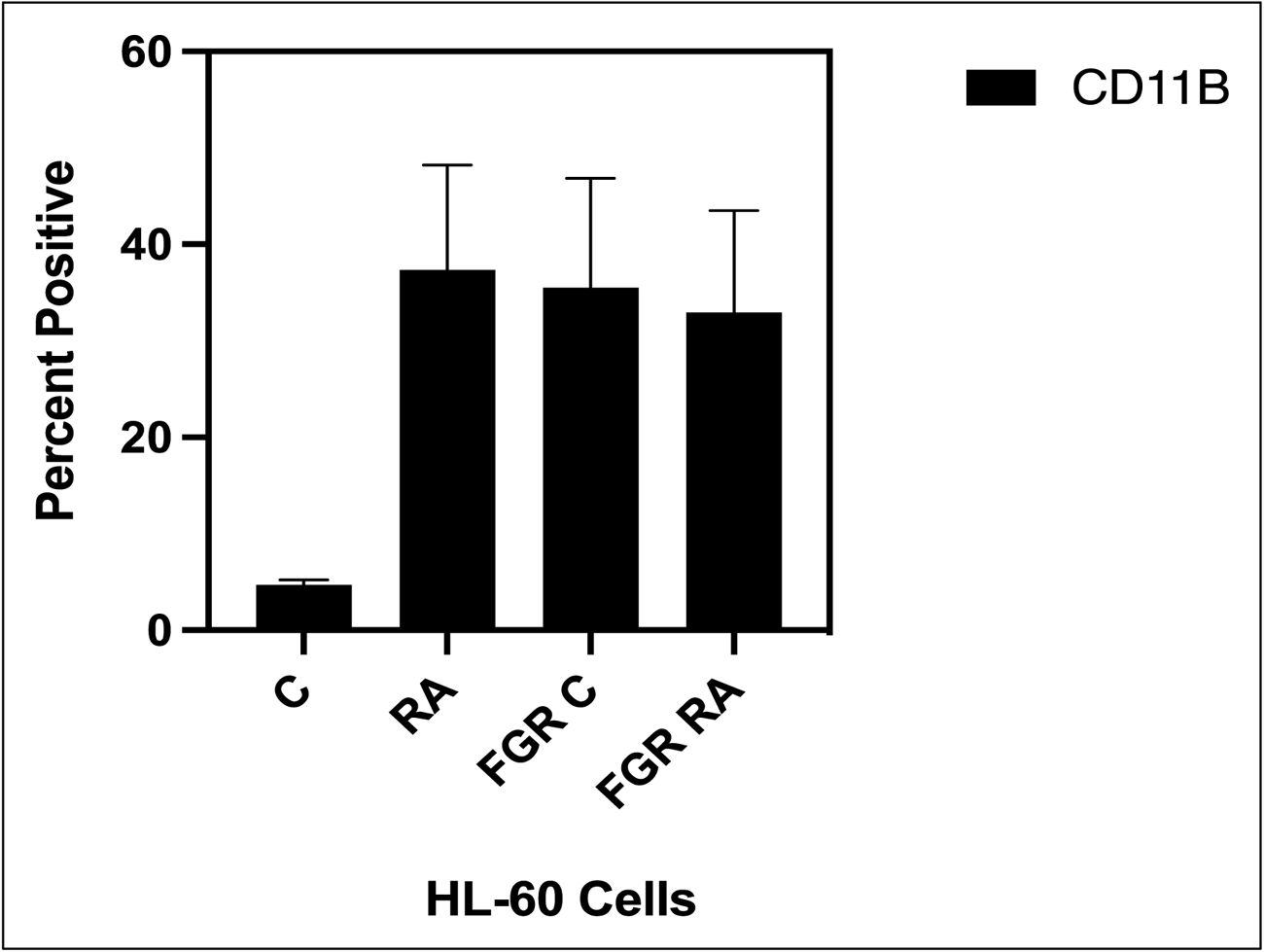
The CD11B expression histogram. Percentage of cells positive for CD11B expression at 72 h (*n* = 3). Error bars indicate SEM comparing RA-treated HL-60 wt samples to RA FGR O.E cells samples.

**Figure S21:**
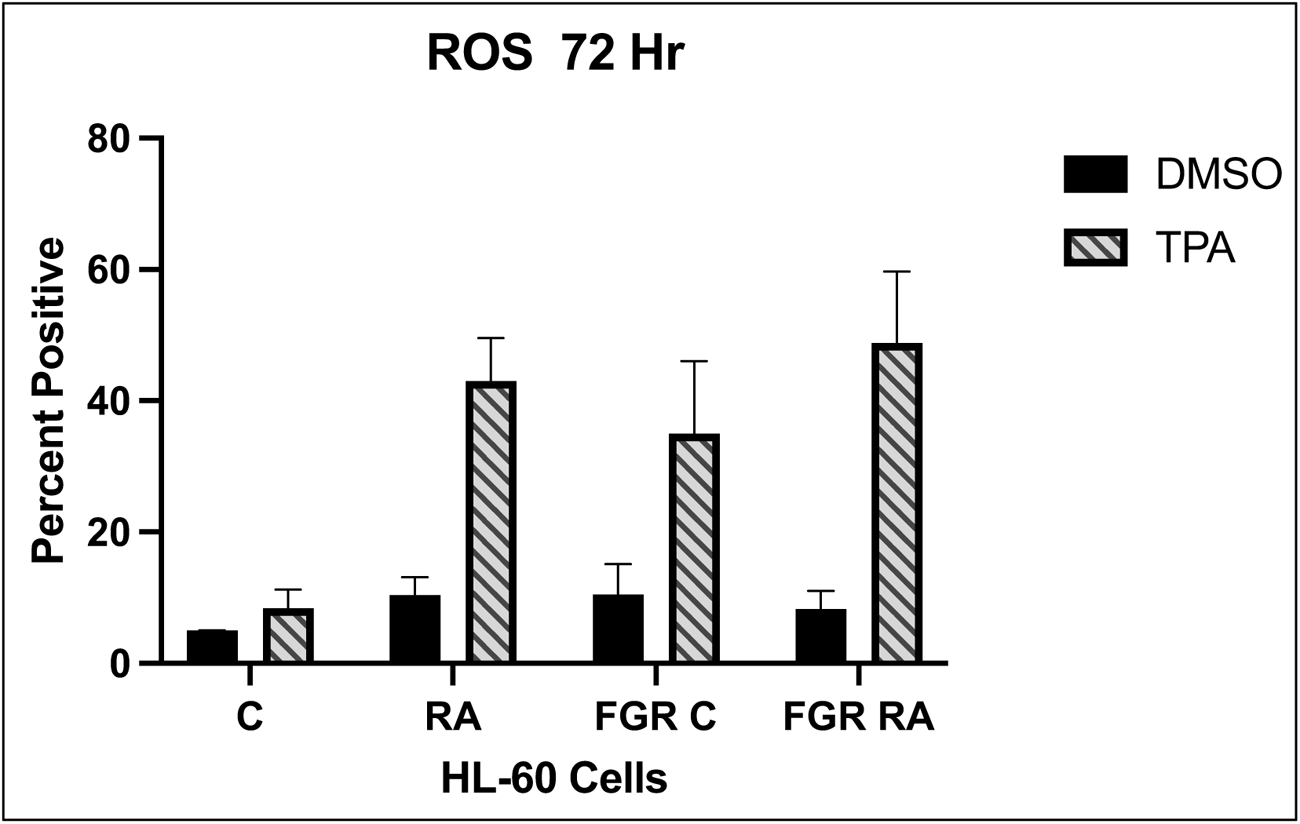
Functional differentiation marker analysis of HL-60 wt and FGR O.E cells untreated and treated with RA measured by TPA-induced respiratory burst. HL-60 WT (parental wildtype) cells were cultured in the absence (control) or presence of 1 μM RA as indicated. FGR O.E cells were cultured in the absence or presence of 1 μM RA as indicated. Respiratory burst was analyzed by measuring reactive oxygen species (ROS) production by flow cytometry using the 2′,7′-dichlorofluorescein (DCF) assay for DMSO carrier control and TPA induced cells. For each of the 4 cases, WT and FGR that were control and RA-treated, TPA-treated samples show induced ROS (*n* = 3). Error bars indicate SEM.

**Figure S22:**
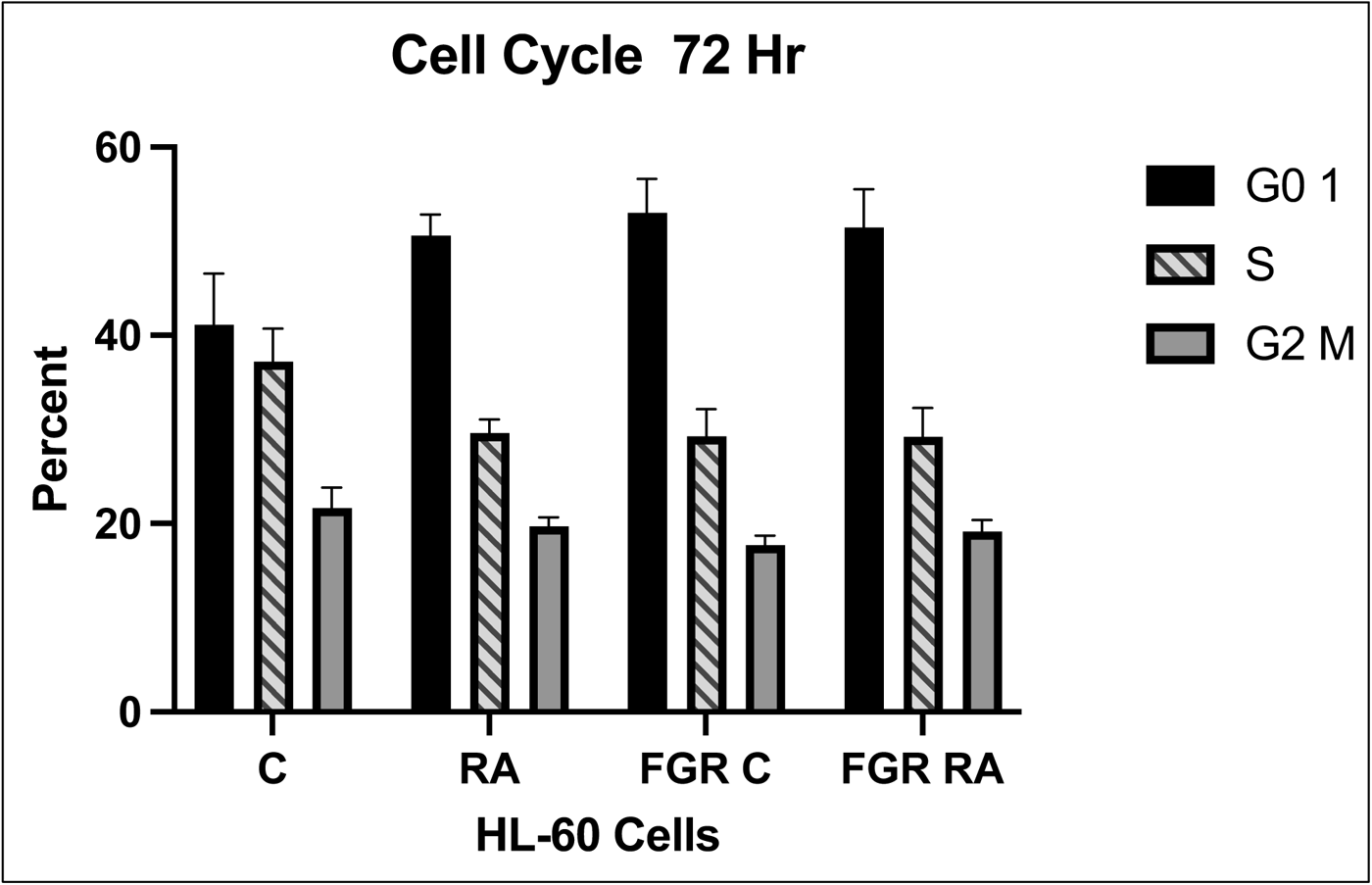
Cell cycle analysis of HL-60 WT and FGR O.E cells. Histogram shows percentage of cells in each phase. Error bars indicate SEM (n = 3) comparing untreated samples to RA/ treated samples.

